# Regulators of ECM Structure Enable Functional Adaptation to Tensile Loading in Tendon Explants

**DOI:** 10.64898/2026.03.05.709185

**Authors:** Emma J. Stowe, Brianne K. Connizzo

**Author notes:** **Correspondence:** Brianne K. Connizzo, Departments of Biomedical and Mechanical Engineering, 44 Cummington Mall, ERB 411b, Boston, MA 02215.

## Abstract

Extracellular matrix (ECM) remodeling is essential for adaptation to changing mechanical demands, yet the mechanisms linking altered strain to functional outcomes remain poorly understood. This study aimed to define molecular and cellular programs driving the adaptation of tendon to increased (exercise) and decreased (disuse) strain. Male murine flexor tendon explants were cultured in incubator-housed tensile bioreactors and subjected to step changes in cyclic strain. After acclimation at 1% cyclic strain, exercise tendons experienced a step increase to 5% strain, while disuse tendons underwent stress deprivation. Increased strain produced significant mechanical adaptations, including increased elastic modulus and failure stress. Multiscale analyses of matrix organization, tissue composition, protein synthesis, signaling factors, and proteolytic activity revealed the mechanisms underlying these adaptations. Exercise-induced functional improvements were linked to an anabolic remodeling program characterized by TGF-β and IL-6 signaling, small leucine-rich proteoglycan expression, MMP suppression, and enhanced collagen alignment. These findings indicate that regulators of matrix organization and turnover, beyond synthesis alone, are critical for functional adaptation. In contrast, mechanical unloading reduced collagen synthesis and alignment and promoted an MMP-dominant, catabolic phenotype favoring matrix breakdown. This study provides a comprehensive characterization of ECM remodeling, linking defined mechanical perturbations to molecular regulation and emergent structure-function relationships. These findings identify targetable mediators of adaptive remodeling and establish a framework for future studies of maladaptive ECM changes in aging, injury, and disease.

## 1 Introduction

Tendons are highly mechanosensitive tissues that transmit force from muscle to bone and enable joint movement. Their function depends on an anisotropic, hierarchically organized extracellular matrix (ECM) composed of longitudinally aligned collagen fiber bundles that confer high tensile strength. While type I collagen is the principal structural protein, non-collagenous matrix constituents, including proteoglycans, glycoproteins, and interstitial water, play critical roles in fibril organization and contribute to the tissue’s nonlinear, viscoelastic mechanical behavior [1]. Importantly, ECM structure is highly dynamic and remodels in response to changes in mechanical stimulation. Defining the mechanisms that regulate ECM remodeling and the resulting structure–function relationships is essential for understanding tendon health, preventing injury, and improving rehabilitation strategies.

Tenocytes sense mechanical forces via cell-cell and cell-ECM interactions and transduce these cues into biochemical signals that regulate ECM remodeling [2,3]. This remodeling process requires tight regulation of matrix degradation and synthesis. Tenocytes secrete proteolytic enzymes, including matrix metalloproteinases (MMPs), that remove damaged matrix, while also producing new matrix proteins that are incorporated into the ECM and reorganized into functional tissue structure. Healthy tendons sense changes in their mechanical environment and can modulate their ECM turnover to meet the functional needs of the tissue [2–4]. It is well established that increased <u>and</u> decreased mechanical loading induces adaptations in tendons [5,6,4]. *In vivo* studies demonstrate that both exercise and disuse alter tendon mechanical properties [7–10] and collagen turnover [11–13]. However, precise control of mechanical stimulation remains challenging even in rodent models, and isolating tendon-specific responses from systemic adaptations and signaling from adjacent tissues (e.g., muscle and bone) is difficult. Tendon explants address these limitations by preserving native cell-cell and cell-ECM interactions while permitting precise mechanical control and enabling real-time assessment of ECM remodeling in an isolated tissue.

Mechanical loading bioreactors have been widely used to model physiologic and pathologic tendon loading *ex vivo* [14–21]. Consistent with *in vivo* observations, stress-deprived tendon explants exhibit a catabolic, MMP-rich phenotype [21–23], while mechanical loading above a threshold promotes anabolic remodeling and suppresses degeneration [16–18,20,21,24]. In a recent study, our group demonstrated strain-dependent regulation of protein turnover and ECM-related gene expression in murine flexor tendon explants subjected to various levels of cyclic tensile strain [14]. However, this study, as well as others in the literature, initiated loading or unloading immediately at the onset of culture, making it challenging to decouple the effects of acute injury and *ex vivo* culture adaptation from true mechanobiological responses. To overcome this limitation, we developed an *ex vivo* model of isolated mechanical perturbations that incorporates a seven-day acclimation period prior to loading interventions, allowing tissues to establish an *ex vivo* baseline state, and isolating the effects of subsequent strain step-changes.

The primary objective of this study was to utilize step changes in tensile strain to uncover molecular and cellular adaptations to both increased (exercise) and decreased (disuse) strain magnitude. We hypothesized that exercise would promote anabolic remodeling characterized by elevated collagen synthesis, preserved matrix organization, and enhanced growth factor signaling, leading to improved mechanical function. Conversely, we predicted that disuse would shift remodeling toward a catabolic state marked by diminished collagen synthesis, disrupted matrix organization, and increased MMP-mediated degradation. By integrating multiscale assessments of ECM turnover, mechanosensitive signaling, and mechanical function, we aimed to link molecular and structural remodeling with functional outcomes to identify key regulators and predictive parameters of tissue adaptation. This work provides mechanistic insight into load-induced tendon adaptation and has potential clinical implications in tendon rehabilitation strategies.

## 2 Results

### 2.1 Adaptive Loading Protocol

We utilized custom-designed, tensile loading bioreactors to subject flexor digitorum longus (FDL) tendon explants to step changes in cyclic strain (Figure 1a). We investigated adaptations to both increased (exercise) and decreased (disuse) strain magnitude. The adaptive loading protocol consisted of distinct acclimation and intervention loading phases (Figure 1b). All groups were first acclimated to *ex vivo* conditions at 1% cyclic strain for 7 days, a physiologic baseline previously established for maintaining FDL tendon explants ex vivo [14]. On day 7, tendons in exercise group underwent a step increase to 5% cyclic strain, while tendons in the disuse group were subjected to stress-deprived conditions for the remainder of the study (Figure 1b). Control tendons were maintained at 1% cyclic strain throughout the culture period.

**Figure 1.**
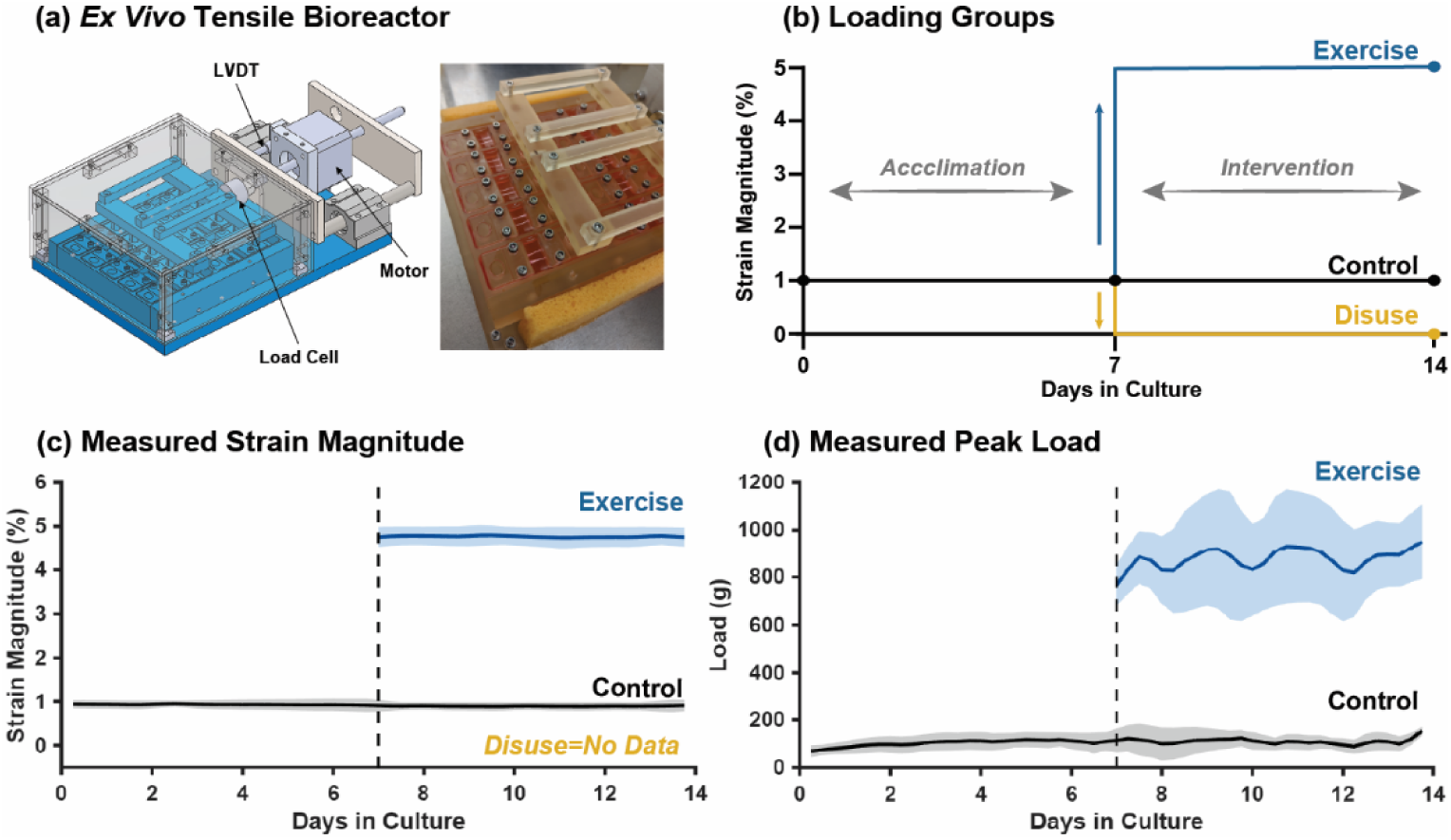
(a) *Ex vivo* tensile-loading bioreactor (left) and experimental setup showing gripped tendon explants (right). (b) Experimental design schematic indicating cyclic strain magnitudes during the acclimation and intervention phases. Three intervention groups were investigated: control, disuse, and exercise. (c) Real-time strain calculated from LVDT displacement readings and (d) peak load measured by the load cell. Solid lines indicate the mean across experiments for each loading condition; shaded areas represent ± one standard deviation. No real-time data are available for the disuse group because it was disconnected from the control system after 7 days.

Real-time load and displacement were measured using a precision load cell and a linear variable differential transformer (LVDT), respectively. Calculated average strain profiles demonstrated high precision and reliable control of the input waveforms (Figure 1c). As expected, average peak loads were elevated during the exercise intervention, although substantial variability was observed between exercise experiments (Figure 1d). Stress-deprived tendons remained gripped but fully slack within the bioreactor system; therefore, real-time load and displacement data were not collected under stress-deprived conditions.

### 2.2 Mechanical Function

To assess functional adaptations of the loading interventions, we quantified both the quasistatic and dynamic mechanical performance of control, disuse, and exercise tendons. Both disuse and exercise groups exhibited decreased cross-sectional area at day 14 compared with control (Figure 2a). Tendon stiffness was highest in the exercise group, showing significant increases compared with control at day 14, and was the only condition to maintain the day 7 reference level (Figure 2b). Elastic modulus was significantly elevated in exercise tendons compared to control, while disuse tendons exhibited an upwards trend in modulus (Figure 2c). Ultimate failure stress was significantly increased only in exercise tendons (Figure 2d). Tissue stress relaxation was assessed at 4%, 6%, and 8% grip strain. At all three strain levels, exercised tendons displayed reduced relaxation compared with control (Figure 2e).

**Figure 2.**
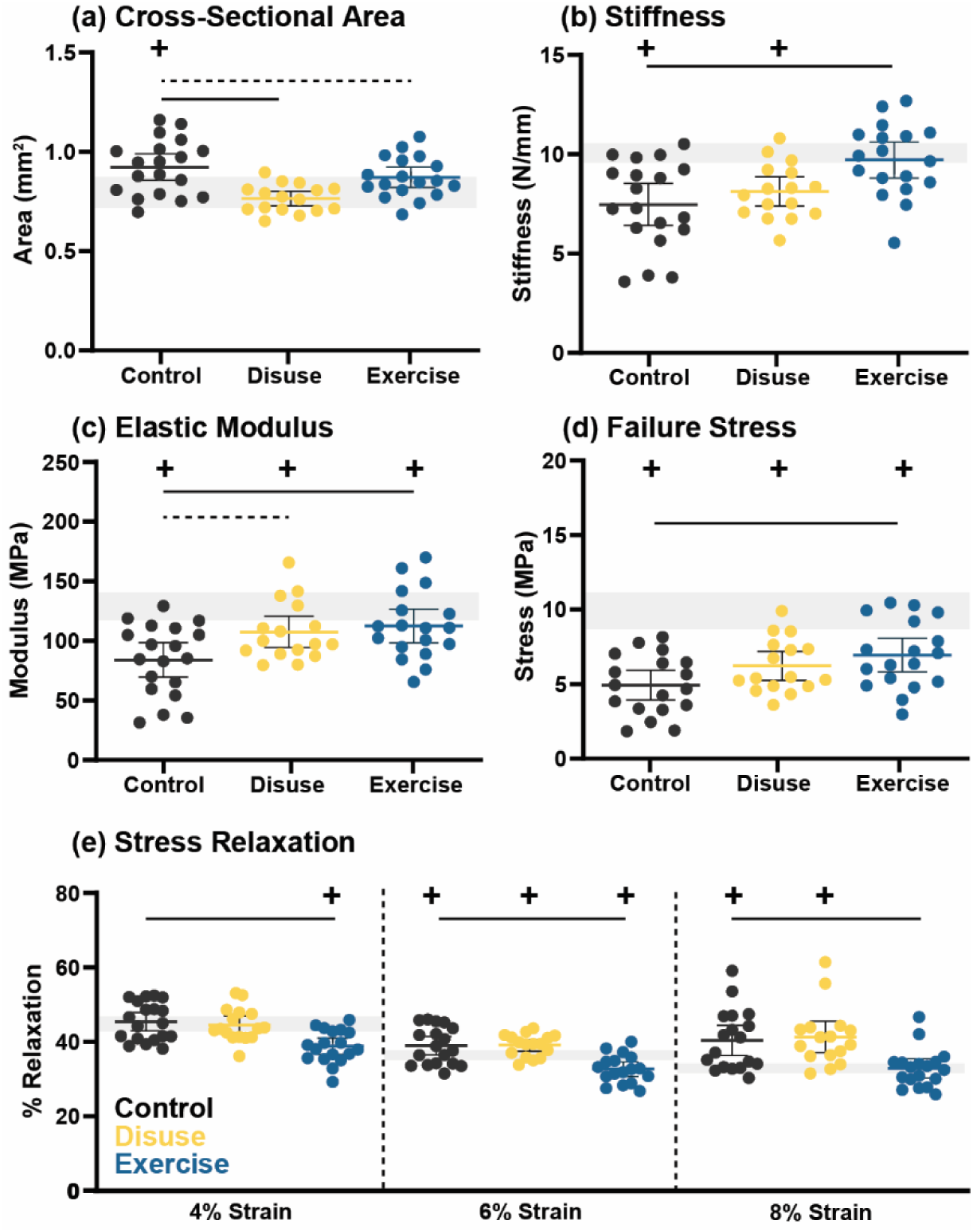
(a) Tendon cross-sectional area, (b) stiffness, (c) elastic modulus, (d) failure stress, and (e) stress relaxation (measured at 4%, 6%, and 8% grip strain) for day 14 control, disuse, and exercise tendons. Data are shown as individual points with mean ± 95% confidence interval. Significant (p < 0.05) comparisons versus control are indicated with a solid line while trends (p < 0.1) are indicated with a dashed line. Grey horizontal bands show the day 7 pre-intervention reference levels (± 95% CI), and (+) symbols indicate significant differences from day 7 (p < 0.05).

Dynamic modulus and tangent of phase angle were determined from frequency sweeps at 0.1 Hz and 1 Hz performed at 4%, 6%, and 8% grip strain. No significant differences were found in either metric at 4% strain (Figure 3a, d). Day 7 dynamic modulus was maintained by both disuse and exercise interventions, but not control, when tested at 6% (Figure 3b) and 8% (Figure 3c) grip strain. At 0.1 Hz, the tangent of the phase angle decreased in exercise tendons compared with control at both 6% (Figure 3e) and 8% (Figure 3f) grip strain.

**Figure 3.**
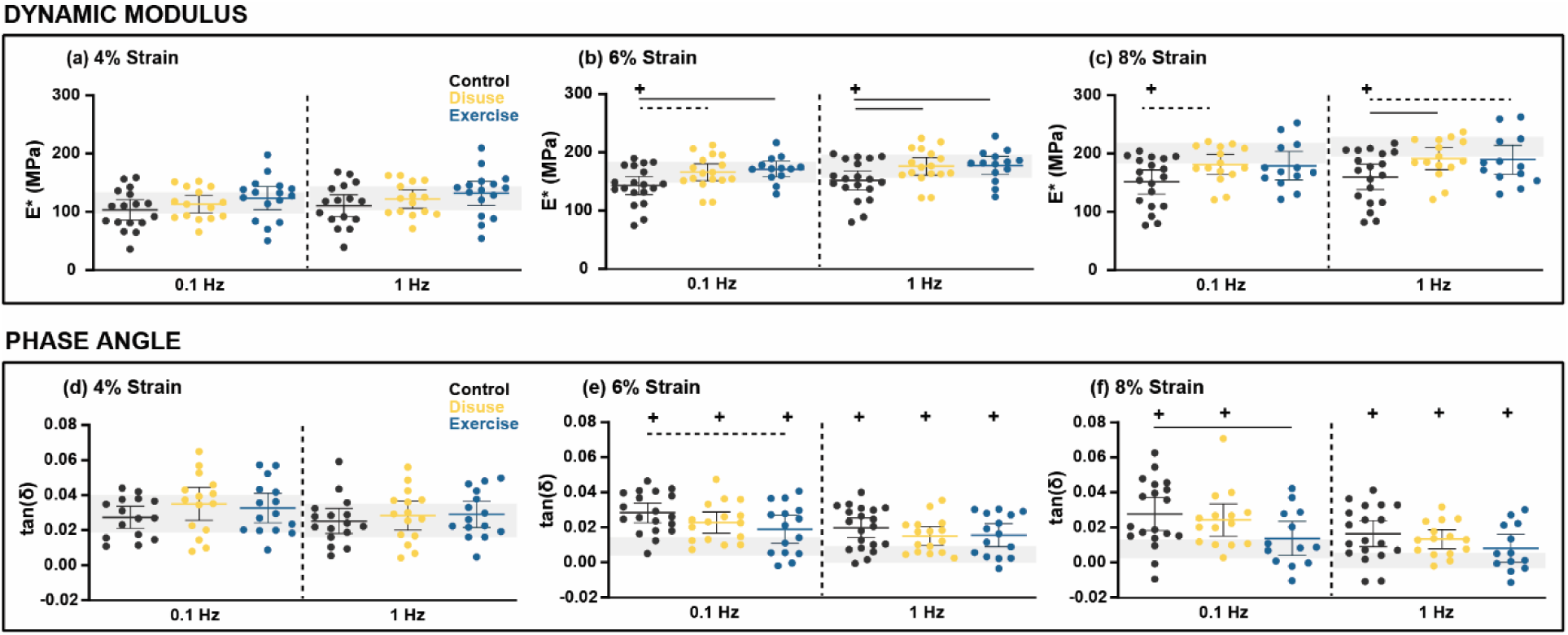
Dynamic modulus (E*, top) and tangent of phase angle (tan(δ), bottom) for day 14 control, disuse, and exercise tendons were calculated from dynamic mechanical testing at 0.1 Hz and 1 Hz. Measurements were performed at (a,d) 4%, (b,e) 6%, and (c,f) 8% grip strain. Data are shown as individual points with mean ± 95% confidence interval. Significant (p < 0.05) comparisons versus control are indicated with a solid line while trends (p < 0.1) are indicated with a dashed line. Grey horizontal bands show the day 7 pre-intervention reference levels (± 95% CI), and (+) symbols indicate significant differences from day 7 (p < 0.05).

### 2.3 Collagen Synthesis and Fibrillar Organization

Seeking to identify mechanisms underlying mechanical adaptations, we assessed collagenous matrix synthesis and fibrillar organization using qPCR, biochemical assays, and second harmonic generation (SHG) imaging. By day 14, all intervention groups showed increased *Col1a1* gene expression, with no differences between groups (Figure 4a). *Col3a1* expression was increased in control and disuse tendons, but not in exercise tendons (Figure 4b). Total protein synthesis, measured by radiolabel incorporation, decreased with disuse and increased with exercise, relative to control (Figure 4c). Additionally, collagen content was decreased in disuse tendons compared to control (Figure 4d). Representative SHG images show collagen structure and organization at day 7 and at day 14 for all three loading interventions (Figure 4g). Quantifications revealed increased collagen fiber dispersion in control and disuse tendons, but not in exercise tendons, between days 7 and 14 (Figure 4e). The collagen alignment index decreased between day 7 and day 14 under control and disuse loading but was maintained with exercise (Figure 4f).

**Figure 4.**
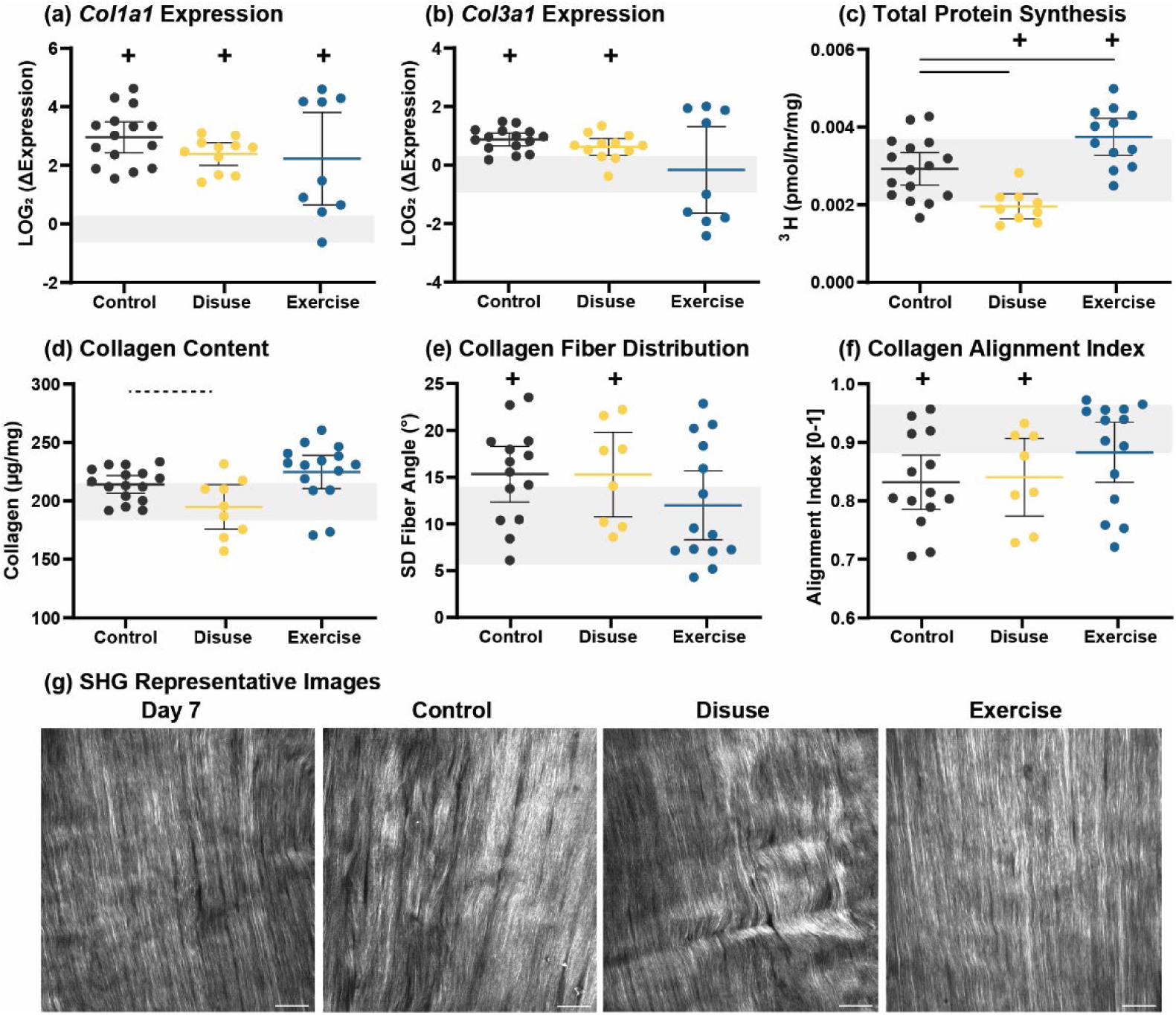
(a) *Col1a1* gene expression, (b) *Col3a1* gene expression (c) total protein synthesis, and(d) total collagen content of day 14 control, disuse, and exercise tendons. (e) Standard deviation of collagen fiber angles (fiber dispersion) and (f) collagen alignment index (AI) determined from SHG images. (g) Representative SHG images with 50 µm scale bar. Data are shown as individual points with mean ± 95% confidence interval. Significant (p < 0.05) comparisons versus control are indicated with a solid line while trends (p < 0.1) are indicated with a dashed line. Gene expression data presented as relative expression to day 7 pre-intervention levels. Grey horizontal bands show the day 7 pre-intervention reference levels (± 95% CI), and (+) symbols indicate significant differences from day 7 (p < 0.05).

### 2.4 Proteoglycan Regulation and GAG Accumulation

To evaluate proteoglycan regulation and non-collagenous matrix composition, we measured expression of the small leucine-rich proteoglycans *Bgn*, *Fmod*, and *Dcn*, sulfated GAG synthesis, total GAG content, and tissue water content in control, disuse, and exercise tendons at day 14. *Bgn* (Figure 5a) and *Fmod* (Figure 5b) expression were significantly increased during the intervention phase for all groups, with no significant differences between groups. Relative to day 7, *Dcn* expression (Figure 5c) was downregulated in control and disuse groups, but unchanged in exercise tendons. All groups exhibited increased sGAG synthesis at day 14 relative to day 7 (Figure 5d). However, total GAG content decreased during the intervention phase for control and exercise tissues; disuse tendon exhibited significantly higher GAG content, maintaining day 7 levels (Figure 5e). Total water content increased in all groups at day 14 relative to day 7 (Figure 5f).

**Figure 5.**
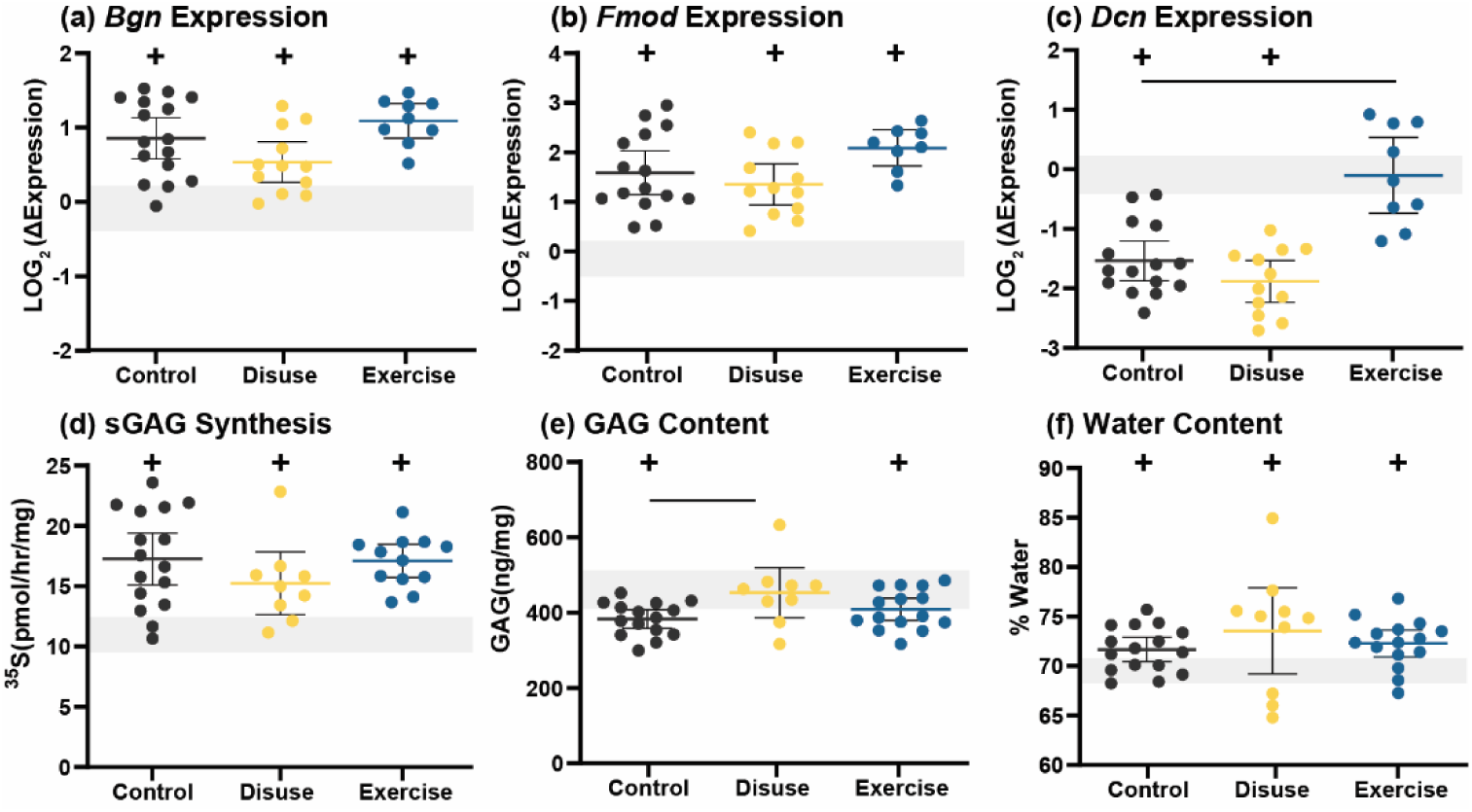
(a) *Bgn* gene expression, (b) *Fmod* gene expression, (c) *Dcn* gene expression, (d) sGAG synthesis, (e) GAG content, and (f) water content of day 14 control, disuse, and exercise tendons. Gene expression data presented as relative expression to day 7 pre-intervention levels. Significant (p < 0.05) comparisons versus control are indicated with a solid line. Data are shown as individual points with mean ± 95% confidence interval. Grey horizontal bands show the day 7 pre-intervention reference levels (± 95% CI), and (+) symbols indicate significant differences from day 7 (p < 0.05).

### 2.5 Mechanosensitive Signaling Factors

To assess mechanically responsive signaling, we measured *Il6*, *Tnfa*, and *Tgfb1* gene expression and quantified fold changes in TGF-β1, TGF-β2, and TGF-β3 protein secretion relative to pre-intervention levels. *Il6* expression was decreased by day 14 in control and disuse tendons but remained unchanged in exercise tendons, yielding a higher relative *Il6* expression in exercise versus control (Figure 6a). By contrast, *Tnfa* expression was unaffected by control and disuse but was downregulated with exercise (Figure 6b). *Tgfb1* expression was upregulated from day 7 to day 14 in all groups; however, relative to control, *Tgfb1* expression decreased with disuse but increased with exercise (Figure 6c). TGF-β1 secretion decreased between day 6 and day 8 in both disuse and exercise groups (Figure 6d). TGF-β2 (Figure 6e) and TGF-β3 (Figure 6f) secretion were significantly upregulated at day 12 in control and disuse conditions but remained unchanged with exercise, resulting in a smaller net increase in TGF-β2 and TGF-β2 release with exercise.

**Figure 6.**
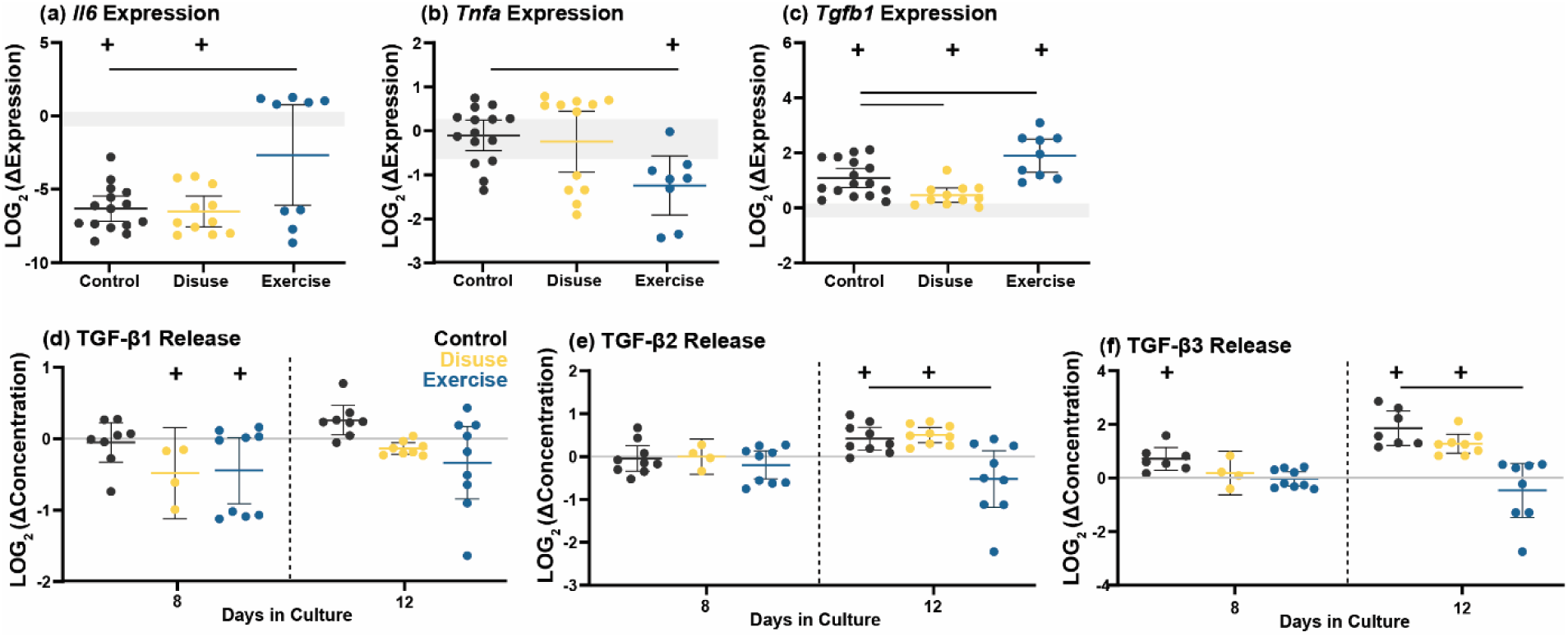
Gene expression of (a) *Il6,* (b) *Tnfa*, and (c) *Tgfb1* in day 14 control, disuse, and exercise tendons relative to day 7 pre-intervention levels. Fold change in (d) TGF-β1, (e) TGF-β2, and (f) TGF-β3 protein concentrations in spent culture medium at day 8 and day 12 relative to day 6 pre-intervention levels. Concentrations are paired values collected longitudinally over the culture period for each experiment. Data are shown as individual points with mean ± 95% confidence interval. Significant (p < 0.05) comparisons versus control are indicated with a solid line. Grey horizontal bands show the day 7 pre-intervention reference levels for gene expression data (± 95% CI), and (+) symbols indicate significant differences from day 6 (protein) or day 7 (gene) (p < 0.05). Raw data in Supporting Figure S6.

### 2.6 MMP Production and Activity

We quantified MMP gene expression, MMP/TIMP protein secretion, and MMP activity to assess matrix degradation. Both *Mmp1* (Figure 7a) and *Mmp13* (Figure 7c) expression showed no differences over the intervention period or between intervention groups. *Mmp3* gene expression was downregulated for all loading conditions between days 7 and 14 (Figure 7b). MMP-3 secretion into culture medium was increased with both disuse and exercise at day 8, and with disuse only at day 12, compared with control (Figure 7d). MMP-9 secretion was upregulated in all groups at both day 8 and day 12; however, the fold increase in MMP-9 release at day 12 was lower in exercise compared with control (Figure 7e). MMP-13 release was upregulated at day 8 and day 12 for control and disuse but not in exercise tendons, resulting in a significantly lower fold change in MMP-13 secretion with exercise (Figure 7f). TIMP-1 secretion was increased in all groups at day 8 relative to day 6; by day 12, exercise tendons produced TIMP-1 levels unchanged from day 6, whereas production remained increased with control and disuse (Figure 7g). Relative MMP activity was significantly increased in disuse tendons compared to control tendons at day 8 (Supporting Figure S5). Raw secreted protein concentrations (not normalized to pre-intervention levels) are provided in Supporting Figure S6.

**Figure 7.**
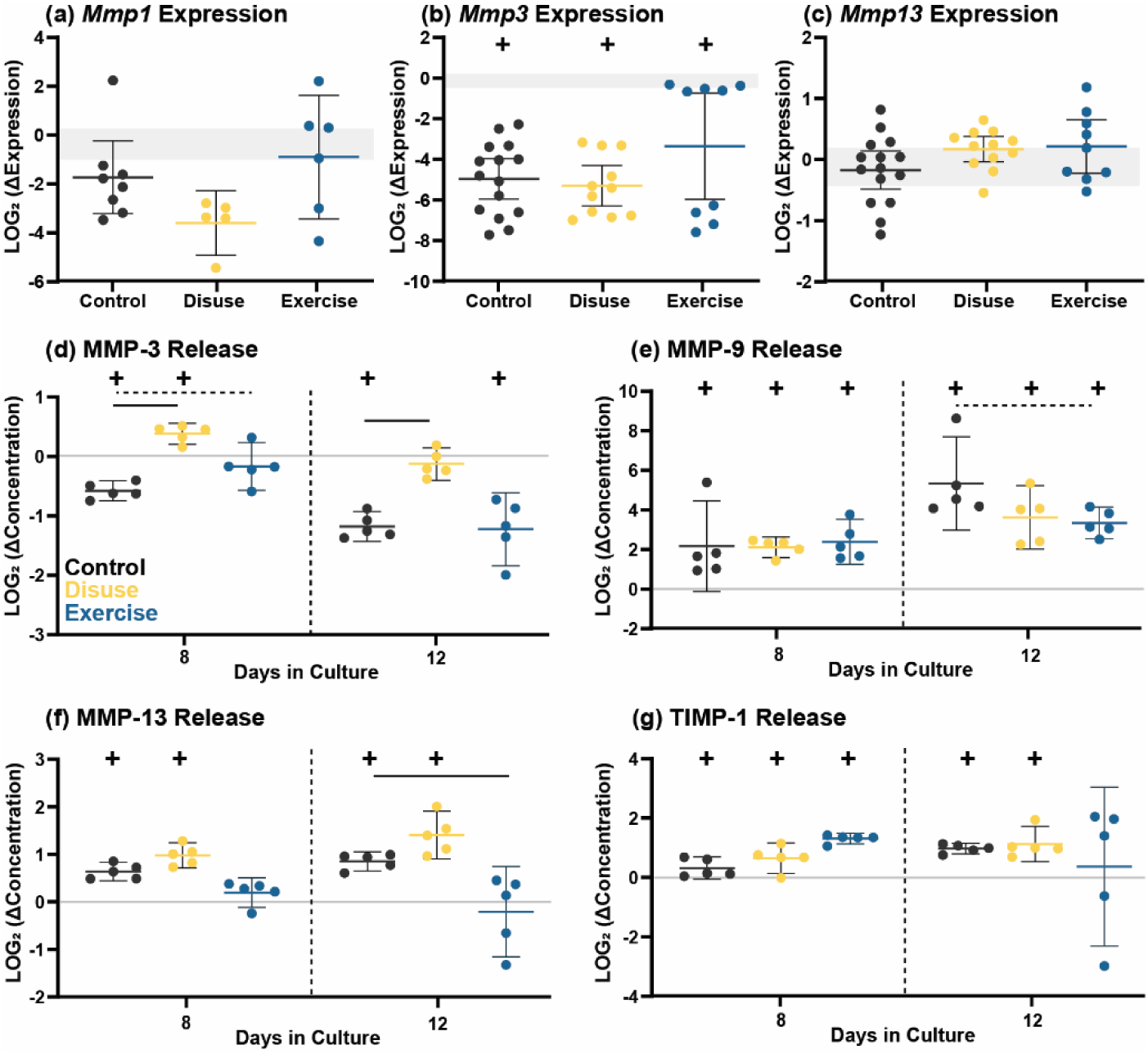
Gene expression of (a) *Mmp1,* (b) *Mmp3*, and (c) *Mmp13* at day 14 in control, disuse, and exercise tendons relative to day 7 pre-intervention levels. Fold change in concentrations of (d) MMP-3, (e) MMP-9, (f) MMP-13, and (g) TIMP-1 in spent culture medium at days 8 and 12 relative to day 6 pre-intervention levels. Concentrations are paired values collected longitudinally over the culture period for each experiment. Data are shown as individual points with mean ± 95% confidence interval. Significant (p < 0.05) comparisons versus control are indicated with a solid line while trends (p < 0.1) are indicated with a dashed line. Grey horizontal bands show the day 7 pre-intervention reference levels for gene expression data (± 95% CI), and (+) symbols indicate significant differences from day 6 (protein) or day 7 (gene) (p < 0.05). Raw data in Supporting Figure S6.

### 2.7 Correlating Mechanistic Adaptations

To integrate multiscale data and link molecular and compositional changes with functional outcomes, we performed a post-hoc correlation analysis on 32 measured variables. Variable abbreviations, units, assays, and timepoints used for the correlation analyses are provided in Supporting Table S2. A heatmap showing the Pearson r correlation coefficients between adaptation parameters is shown in Figure 8a. To examine these relationships further, we performed simple linear regressions between selected variable pairs (Figure 8b-i). Color-coding the pooled data by loading intervention allows visualization of both the parameter relationships and the effects of different mechanical states on data distribution. Elastic modulus showed a moderate positive correlation with collagen alignment (Figure 8b) and moderate negative correlations with release of TGF-β1, TGF-β2, TGF-β3, MMP-13, and TIMP-1 (Figure 8a, c). Conversely, tendon stress relaxation demonstrated a negative relationship with collagen alignment (Figure 8d) and positive relationships with the release of TGF-β and MMP family proteins (Figure 8a, e). Total collagen content showed a moderate positive correlation with proline incorporation (Figure 8f) and a moderate negative correlation with MMP-3 and MMP-13 release (Figure 8a, g). Interestingly, TIMP-1 release showed a strong positive correlation with total collagen content (Figure 8a; individual regression not shown). Collagen alignment showed moderate-to-strong negative correlations with the release of TGF-β and MMP family proteins (Figure 8a; individual regressions not shown). Total GAG content showed a very weak positive correlation with sulfate incorporation (Figure 8h) but a strong positive correlation with MMP-3 release (Figure 8i).

**Figure 8.**
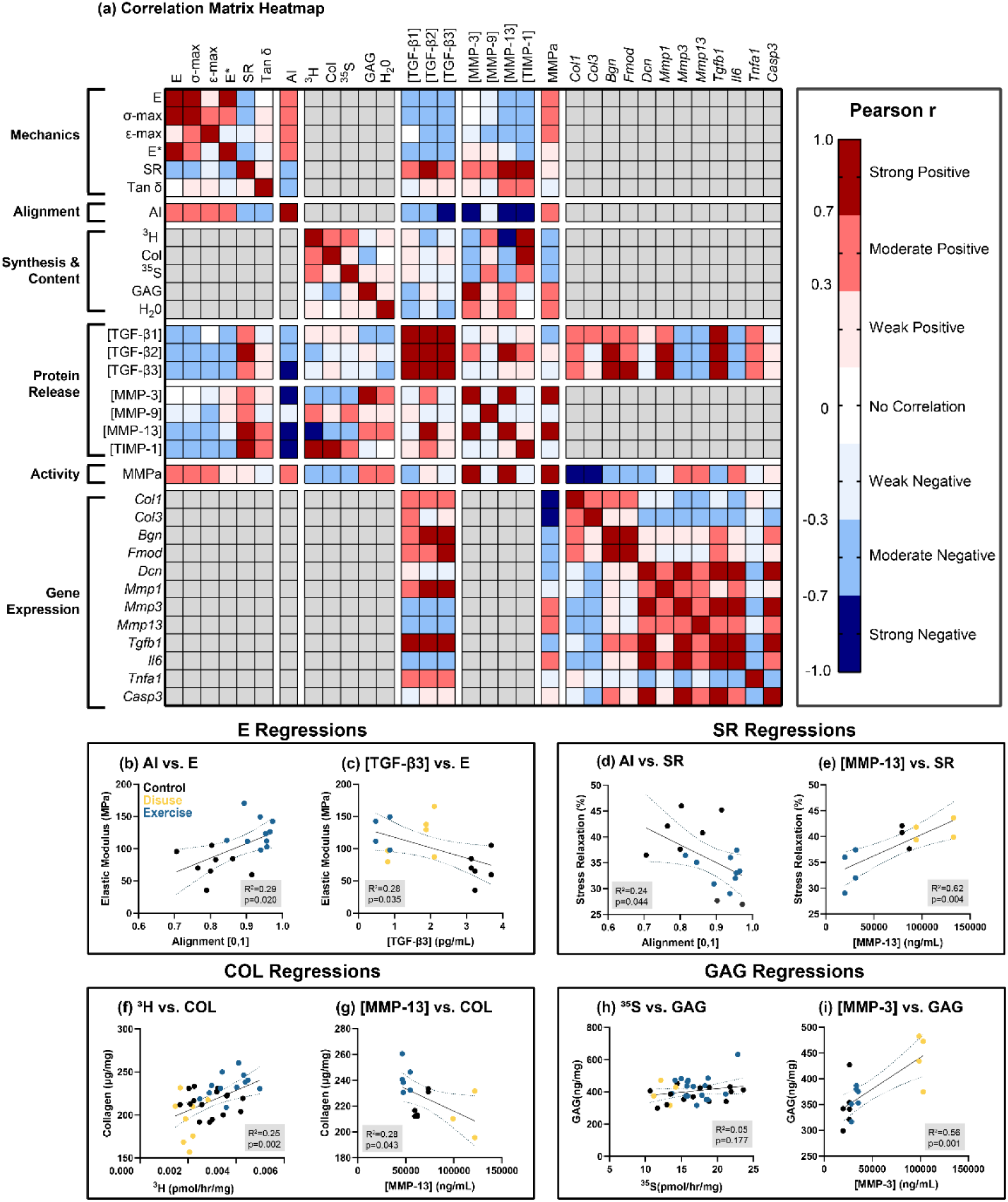
(a) Heatmap showing the Pearson correlation matrix (r) for adaptation parameters measured in individual tendon explants. The legend (right) indicates seven color levels ranging from strong negative correlation (dark blue, −1 ≤ r ≤ −0.7) to strong positive correlation (dark red, 0.7 ≤ r ≤ 1). Parameters with no correlation (r = 0) are shown in white, and parameters lacking sufficient data are shown in grey. Select simple linear regressions are shown for (b,c) elastic modulus (E), (d,e) stress relaxation (SR), (f,g) collagen content (COL), and (h,i) glycosaminoglycan content (GAG). Correlations and regressions include data pooled across all loading interventions. Panels (b–i) show individual data points colored by loading intervention, with regression lines and 95% confidence intervals. R² values and p-values for slope significance are shown in inset grey boxes. Variable abbreviations (Table S2), correlation coefficients (Table S3), p-values (Table S4), and sample sizes (Table S5) are provided in the Supporting Information.

## 3 Discussion

ECM remodeling is central in the ability of tendons to respond to changing mechanical demands. Failure to adapt, as occurs with aging or disease, compromises tissue integrity and increases risk of injury. Thus, defining the mechanisms that regulate ECM adaptation is critical for predicting and modulating functional outcomes associated with dysregulated remodeling. In this study, we examined ECM remodeling in young male tendon explants subjected to discrete step changes in tensile strain magnitude. We found that increased strain produced significant adaptations in both elastic and viscoelastic mechanical properties, including increased elastic stiffness, higher failure resistance, and reduced viscous energy dissipation, consistent with prior *in vivo* exercise studies in rodents [13] and humans [10,11,33–35]. We then aimed to link these functional outcomes with molecular and structural adaptations via multiscale analyses of matrix organization, tissue composition, protein biosynthesis, secreted signaling factors, and enzymatic activity. Our findings demonstrate that tendon extracellular matrix remodeling is governed primarily by regulators of collagenous matrix turnover and organization, rather than synthesis alone. Specifically, exercise-induced improvements in mechanical function were most strongly associated with an anabolic remodeling program characterized by TGF-β and IL-6 signaling, small leucine-rich proteoglycan (SLRP) expression, MMP suppression, and collagen organization.

It is well established that tendon mechanical behavior is highly dependent on matrix organization [33–35]. Consistent with this relationship, we observed preservation of collagen organization in exercised tendons and significant correlations between collagen alignment and mechanical properties (Figure 8). While these results emphasize that matrix organization is necessary for improved mechanical function, productive remodeling is also highly dependent on the capacity to incorporate newly synthesized proteins, as well as the magnitude and type of collagen produced. Proline incorporation, a widely accepted corollary for newly synthesized and incorporated collagen [3], increased with exercise and decreased with disuse (Figure 4c). Previous *in vivo* studies have reported similar increases in collagen synthesis following both acute and longer-term exercise in human Achilles and patellar tendons, corroborating our results [11,12,36]. Comparable findings have been reported *ex vivo*, with 24-hours of 5% cyclic strain increasing proline incorporation in isolated rat tendon fascicles [16]. We also found a positive correlation between collagen synthesis and total collagen content, confirming that higher synthesis rates are associated with net collagen accumulation (Figure 8f). In contrast, disuse tendons exhibited decreased collagen content and synthesis, supporting a degenerative phenotype with mechanical unloading (Figure 4c, d). While *Col1a1* expression increased in all groups during the intervention, *Col3a1* was upregulated only in control and disuse tendons (Figure 4a, b). Wang *et al*. reported similar results in an *ex vivo* rabbit explant model, showing that stress deprivation increased collagen III type gene expression and immunofluorescence while cyclic loading restored levels toward those of native tendon [17]. As collagen III type is associated with immature or provisional ECM and collagen type I reflects mature, load-bearing matrix [37], reduced *Col3a1* with exercise could reflect a shift from initial neomatrix production toward a collagen I–dominant, mechanically optimized matrix. These observations support a model in which increased cyclic strain promotes anabolic remodeling, enhancing collagen synthesis and fiber alignment to improve elastic function.

Importantly, we demonstrate that tendon adaptation is governed by regulation of collagenous matrix turnover and organization, rather than by increased matrix synthesis alone. Collagenous matrix organization is tightly regulated by non-collagenous matrix components that control fibrillogenesis and collagen assembly, shaping tendon mechanical behavior [1]. Small leucine-rich proteoglycans (SLRPs) such as decorin, biglycan, and fibromodulin are central to fibril formation and matrix assembly, and SLRP dysregulation has been reported in settings of altered matrix turnover and tendon pathology [31,38–43]. Decorin expression was maintained in exercised tendons but decreased in control and disuse groups (Figure 5c), yielding significantly higher decorin expression at day 14 in exercised tissues. Coupled with improved collagen fiber alignment, these data support a mechanistic role for decorin in maintenance of collagen organization even during adulthood. Consistent with our findings, Robinson *et al*. observed altered fibril structure and impaired mechanics in an inducible knockout of decorin and biglycan in mature tendons [39]. Similarly, Xu *et al*. reported increased decorin expression in rat Achilles tendons following moderate treadmill running [44]. Proteoglycans consist of a core protein bound to one or more glycosaminoglycan (GAG) side chains, known for their negative charge and hydrophilicity, which impact tissue hydration and viscoelastic behavior [45]. Consistent with prior observations by Screen *et al*., cyclic loading did not alter sulfated GAG (sGAG) synthesis, despite inducing significant loading-dependent changes in collagen synthesis [16]. However, disuse tendons did exhibit the highest total GAG content at day 14 (Figure 5d–f), a finding consistent with our prior work showing increased GAG content in stress-deprived tendons versus cyclically loaded tendons [14]. We did not detect a significant relationship between sGAG incorporation and total tissue GAG content (Figure 8h), indicating differences in GAG retention or turnover that could affect tissue mechanics and remodeling capacity. One possibility is that newly incorporated sGAG represents a transient pool that participates in short-term matrix regulation and is rapidly turned over. Mechanisms that could underlie differential GAG retention include changes in proteoglycan processing and altered proteolytic balance (MMP/TIMP activity), which we address below.

Matrix metalloproteinases (MMPs) are principal mediators of matrix degradation and core regulators of ECM turnover. The balance between MMPs and their inhibitors (TIMPs) can determine whether the ECM undergoes adaptive remodeling, stable maintenance, or pathological degeneration. Although MMP gene expression was largely unchanged across loading interventions, secreted MMP proteins exhibited load-dependent patterns that paralleled functional outcomes. The exercise group exhibited a late-stage reduction in both MMP-13 and MMP-9 release (Figure 7e, f). As MMP-13 is the principal collagenase for fibrillar collagen and MMP-9 degrades collagen type IV and denatured collagens, these findings are consistent with preservation of the collagenous matrix. Supporting the role of MMPs in matrix breakdown and mechanical decline, we observed significant negative correlations between MMP release and both collagen content and elastic modulus (Figure 8a). Similar observations were reported by Rooney *et al*., who found reduced MMP activity accompanied improved tissue mechanics in rat supraspinatus tendons following 8-weeks of treadmill exercise [13]. Interestingly, during exercise TIMP-1 secretion showed an early transient increase with a return toward baseline by day 14, consistent with temporally controlled regulation of proteolytic activity. Moreover, we found positive correlations between TIMP-1 concentration and collagen content (Figure 8a), underscoring the importance of TIMPs in modulating MMP activity and protecting matrix integrity. Future studies will investigate TIMP transcriptional regulation and assess the production of additional TIMP family members.

Notably, within 24 hours of unloading we detected increased MMP-3 release (Figure 7d) and elevated generic MMP activity (Supporting Figure S5), indicating a rapid proteolytic response to loss of tensile load. This high sensitivity to unloading suggests MMP activation is an early driver of matrix degradation during the disruption of mechanical homeostasis. Prior stress-deprivation studies corroborate these findings, reporting rapid upregulation (< 24 hours) of MMP-1, MMP-3, MMP-9, and MMP-13 in rat Achilles, supraspinatus, and mouse/rat tail tendon models [18,23,46–48]. Importantly, the upregulated MMP phenotype can be reversed by static or cyclic loading [18,23,46,47,49] and pharmacologic inhibition of MMPs has been shown to prevent loss of mechanical properties during stress deprivation [50]. Together with the exercise data, our findings suggest that MMP-3, a stromelysin that degrades proteoglycans and laminin, is critical for acute remodeling after loss of mechanical stimulus, whereas MMP-9 and MMP-13 appear to act as later responders involved in longer-term regulation of homeostatic remodeling. Interestingly, we observed strong positive correlations between generic MMP activity (MMPs -1, -2, -3, -7, -8, -9, -10, -13, and -14) and the production of MMP-3 and MMP-13 (Figure 8a), implicating MMP-3 and MMP-13 as major contributors to the net proteolytic response in tendon. Given that not every individual MMP was directly quantified here, future work will expand protease profiling and test the specific roles of individual MMPs via targeted supplementation or inhibition.

TGF-β is a signaling molecule strongly associated with mechanical loading and has been implicated as a key regulator of load-induced collagen synthesis both *in vivo* and *in vitro* [51–55]. We assessed TGF-β isoforms and other mechanically-sensitive signaling molecules to investigate pathways implicated in tendon adaptation to mechanical load in our tendon explant model. *Tgfb1* gene expression revealed distinct strain-dependent adaptations with exercise upregulating and disuse downregulating the relative expression (Figure 6c). While these trends did not translate to the protein level, we did observe strain-, time-, and isoform-dependent TGF-β dynamics. TGF-β1 release was transiently decreased in exercise and disuse tissues at day 8 but returned to baseline at day 12 (Figure 6d). TGF-β2 and TGF-β3 exhibited minimal changes at day 8 but showed significantly increased release at day 12 in control and disuse tissues, resulting in a lower net TGF-β2/3 release with exercise (Figure 6e, f). This apparent discrepancy may be explained by TGF-β–ECM binding dynamics [56–58]. TGF-β is synthesized and secreted predominantly in a latent form that is rapidly sequestered within the extracellular matrix, requiring mechanical or proteolytic release to generate the active ligand capable of receptor binding. As a result, TGF-β levels measured in the culture supernatant may not accurately reflect the pool of matrix-bound, activatable TGF-β within the tissue. The increased TGF-β2/3 release observed in control and disuse tendons may therefore reflect greater matrix turnover or proteolytic activity, whereas reduced release with exercise may indicate localized retention of latent TGF-β. Future work will quantify both latent and active pools of TGF-β within the tendon matrix and will assess downstream Smad signaling to directly evaluate TGF-β pathway activation.

We also observed strain-dependent gene expression of *Il6* and *Tnfa* with *Il6* expression increasing and *Tnfa* expression decreasing with exercise (Figure 6 a,b). Prior studies have reported robust IL-6 responses to mechanical loading, with increased IL-6 gene expression in bovine tendon fascicles and in human tendon fibroblasts following *in vitro* cyclic loading [59,60]. In a randomized controlled trial, systemic IL-6 infusion after treadmill running increased tendon collagen synthesis markers, supporting a role for IL-6 in promoting collagen production [61]. It is hypothesized that IL-6 and TGF-β signaling act synergistically to influence matrix synthesis and remodeling [62,63], although the precise mechanisms and context dependence of this crosstalk remain to be defined. Mechanosensitivity of TNF-α, commonly associated with injury and inflammation, is far less documented in the literature. The downregulation of *Tnfa* with exercise that was observed could reflect an anti-inflammatory shift associated with a pro-remodeling phenotype [64].

One of the primary goals of this study was to propose a regulatory framework for how tendons adapt to changing mechanical loads (Figure 9). Based on our findings, we propose a mechanism in which increased tensile loading activates mechanosensitive signaling pathways (TGF-β and IL-6) that promote collagen synthesis. In parallel, preservation of small leucine-rich proteoglycans (SLRPs) supports fibrillogenesis and collagen fiber alignment. The combined effect of increased synthesis and improved organization, together with loading-induced reduction in MMP proteolytic activity, preserves the collagenous matrix and improves tendon elastic function. Conversely, mechanical unloading shifts the balance toward reduced mechanosensitive signaling, decreased collagen synthesis and alignment, and an MMP-dominant phenotype that favors breakdown of disorganized matrix. Functionally, these data support the canonical hypothesis that increased loading favors net matrix production and architecture conducive to load transmission, whereas decreased loading favors net degradation and mechanical weakening.

**Figure 9.**
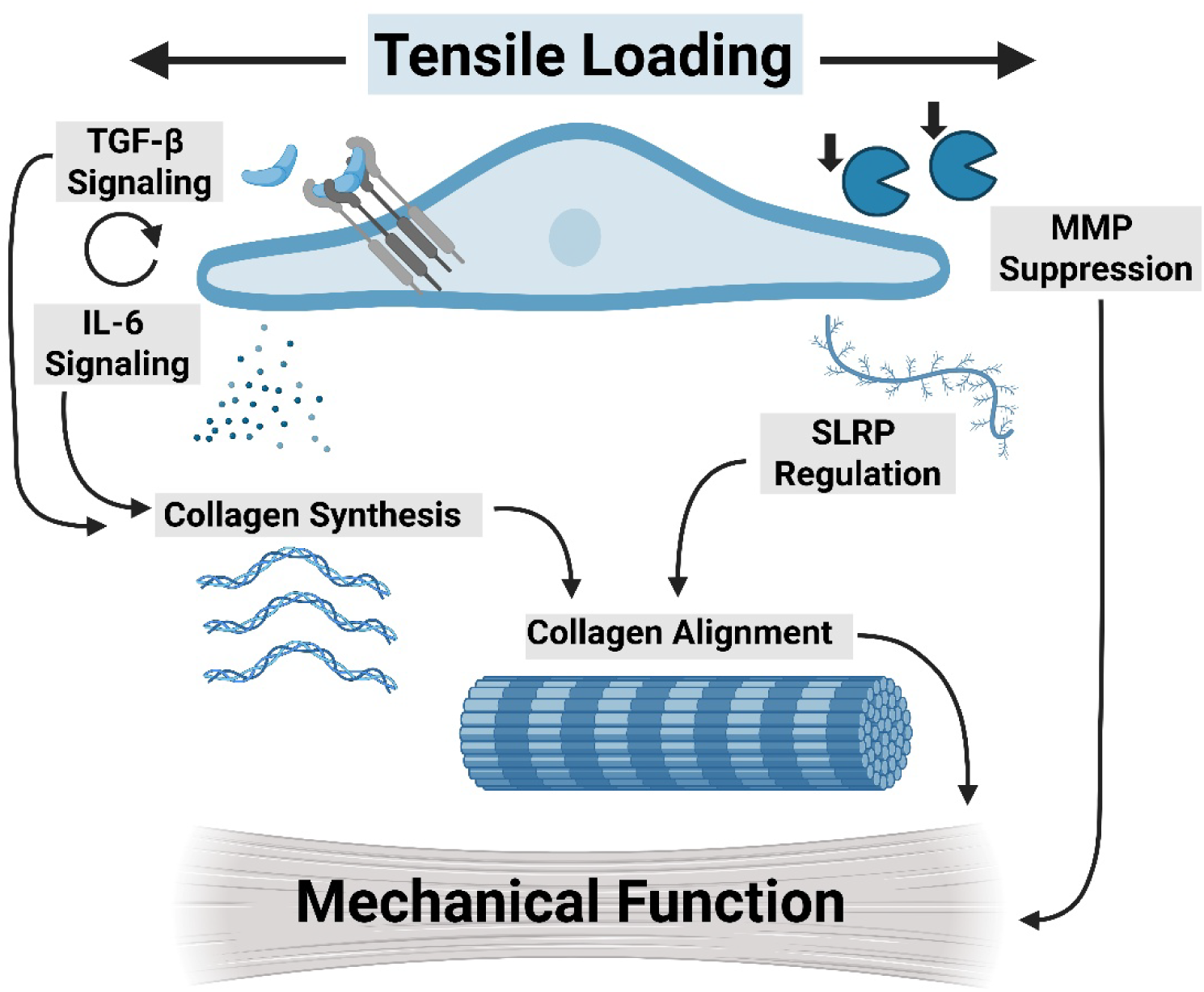
Proposed mechanism of mechanically driven tendon extracellular matrix remodeling. Tensile loading activates mechanosensitive signaling (TGF-β and IL-6) and promotes SLRP regulation, promoting collagen synthesis and fibrillogenesis. Improved collagen organization and reduced proteolytic pressure (MMP suppression), preserves the collagenous matrix and enhances mechanical function.

Beyond this descriptive model, our results raise several mechanistic and translational implications. First, the concurrent regulation of TGF-β/IL-6 signaling and SLRP expression suggests that loading not only tunes synthetic output but also governs the quality and organization of newly produced ECM, implying that therapeutic strategies that stimulate synthesis without restoring proper fibrillogenesis may yield mechanically inferior tissue. Additionally, balanced regulation of MMP-mediated degradation is critical for downstream remodeling, as excessive activity promotes matrix breakdown, and insufficient proteolysis may contribute to fibrotic matrix accumulation. Future studies should manipulate these regulators individually (e.g., SLRP overexpression to promote fibril assembly, selective MMP inhibition to reduce breakdown, or timed TGF-β activation to enhance synthesis) to establish which interventions restore organization and rescue mechanics. Our data also suggest a strain-sensitive window for tendon remodeling. Modest increases in tensile strain improved organizational metrics and mechanics, whereas the absence of load produced a divergent, degradation-dominated response. Defining this window *in vivo* (strain magnitude, duration, and temporal sequence of signaling changes) will be critical to translate these findings into load-based rehabilitation protocols. In clinical cases that necessitate tendon unloading (e.g., immobilization or prolonged bed rest), defining the optimal timing to reintroduce to mechanical loading could have substantial impact on improving tendon healing and recovery. Finally, by linking molecular signaling to matrix architecture and whole-tissue mechanics, our framework generates testable hypotheses for biomaterial and cell-based therapies in which scaffolds or growth factor regimens can couple pro-synthetic signaling with structural cues that guide optimal collagen organization and restore native tendon function.

In addition to mechanistic conclusions about load-driven remodeling, our results have important methodological implications for tendon explant research. We demonstrate that tendon explants remain mechanically sensitive for up to 14 days in *ex vivo* culture, validating explant models for investigations that require multi-week tissue culture and mechanobiological interrogation [65]. Using a novel adaptive loading protocol, explants were acclimated under 1% cyclic strain for 7 days, then subjected to 7 days of loading interventions. The baseline 1% strain was selected based on our prior work demonstrating that 1% strain best maintains native tendon physiology during *ex vivo* culture [14]. Compared with higher strain magnitudes (3%, 5%, or 7%), 1% strain preserved native tendon gene expression, ECM synthesis, metabolic activity, and MMP activity over 7 days [14]. Nevertheless, in this current study, time-dependent changes were still evident in the 1% control group between days 7 and 14 (Supporting Figures S3–S4), characterized primarily by a decline in markers of native matrix maintenance and a relative increase in measures associated with tissue degradation. In contrast, higher strain appeared necessary to preserve the baseline phenotype at later time points. Exercise loading (5% cyclic strain) better maintained collagen alignment as well as *Col3a1* and *Dcn* expression compared with continuous 1% loading. This pattern suggests that higher tensile stimuli may be required to sustain tissue integrity and prevent a proteolytic shift toward matrix breakdown during extended *ex vivo* culture.

Comparing the strain-dependent ECM adaptations here with our prior non-acclimated study indicates that mechanical acclimation unmasks specific mechanobiological responses. For instance, whereas no difference in collagen synthesis was detected between 1% and 5% strain in our prior non-acclimated study [14], acclimated explants in the current study showed a significant increase in collagen synthesis at 5% cyclic strain. This shift likely reflects recovery from explant harvest associated stress and re-establishment of cell–matrix homeostasis, which together restore load sensitivity. Key stimulus parameters for exercise-induced ECM remodeling include loading magnitude, frequency, duration, and mode [4]. *In vivo* measurements of the human Achilles tendon report mean strains of ∼3.1% during walking and ∼6.5–7.9% during running, with peak strains up to ∼13.7% during high-intensity activities, underscoring the wide physiological range of tendon loading [66]. In this study, we selected 5% cyclic strain as our exercise condition based on prior data in murine tendons demonstrating that this magnitude induces significant ECM remodeling without eliciting an injury-associated phenotype (Supporting Figure S2).

Stress deprivation is a common mechanical state used in tendon explant culture to indirectly perturb the effects of mechanical loading on tendon function [17,18,20,21,25,47,67]. A hallmark of these studies is loss of mechanical function with stress deprivation, typically accompanied by upregulation of MMP expression [17,18,21,22,47]. Contrary to our initial hypothesis, we did not observe a significant decline in mechanical function after 7 days of mechanical unloading; instead, disuse tendons showed modest upward trends in elastic and dynamic modulus. Although unexpected, similar findings have been reported in an *in vivo* Botox model of Achilles unloading [68], and may reflect increases in cross-sectional area associated with GAG accumulation and tissue swelling that can mask unchanged material stiffness. In addition, our seven-day acclimation period may have reduced the acute degenerative burden experienced upon culture onset, thereby better preserving mechanical properties and highlighting the importance of strain history in determining remodeling outcomes. Despite preserved or slightly increased mechanical measures, disuse tendons did exhibit several degenerative signatures, reduced collagen synthesis, increased MMP activity, and elevated GAG content, supporting the interpretation that unloading promotes a degradative state.

Several limitations to this study warrant discussion. Most notably, substantial bioreactor-to-bioreactor variability was observed. This study encompasses data from 25 independent bioreactor experiments, with each experiment yielding samples for a single loading condition or time point. The technical complexity of these experiments highlights the challenges associated with achieving precise and reproducible high-throughput mechanical loading. These findings underscore the importance of rigorous load and displacement validation for each experiment, as well as the need for real-time signal feedback during dynamic loading protocols. To mitigate experimental variability, data were pooled from three or more validated experiments per condition; nevertheless, we attribute a significant portion of the observed variability to these technical limitations. Another limitation is that we primarily assessed ECM adaptation at a single timepoint (day 14) which represents a late-stage snapshot of remodeling and does not capture the temporal dynamics or distinct phases of load-dependent ECM adaptation. Ongoing work in our laboratory is focused on the development of a longitudinal bioreactor platform that will enable repeated assessment of load-driven tendon remodeling within the same sample across multiple time points. Additionally, we would like to discuss our chosen terminology, referring to increased strain-magnitude as “exercise” throughout this study. True exercise adaptations *in vivo* involve complex systemic contributions from cardiovascular, neurological, and musculoskeletal systems [69]. In the absence of these systemic mediators, the explant model relies exclusively on local mechanotransduction; thus, the observed “exercise” phenotype should be interpreted as intrinsic responses to mechanical loading rather than integrated organismal adaptations. Finally, several limitations apply to the post hoc correlation analysis. Due to technical constraints, not all variables were measured simultaneously, resulting in gaps within the correlation matrix (Figure 8a). Additionally, due to our experimental design, each media sample reflects protein secretion from two “double-loaded” tendons. To enable correlation with single-tendon endpoint measurements, media concentrations were divided by two, which provides only an approximation of protein production at the individual tendon level.

This investigation establishes a framework for future studies aimed at identifying the mechanistic regulators of load-dependent ECM turnover. Future work will assess the maturity and functional integrity of the collagen network by quantifying lysyl oxidase (LOX)–mediated enzymatic crosslinking, a mechanosensitive process central to collagen fibril stabilization and key determinant of tendon mechanical properties [6,70,71]. While the present study identifies correlations between molecular regulators and functional outcomes, interventional approaches are necessary required to establish causality within this regulatory framework. Specifically, targeted upregulation, downregulation, or inhibition of key regulators (TGF-β, IL-6, and decorin) will enable direct testing of mechanistic links between regulatory signaling, collagen synthesis, matrix organization, and mechanical function. Finally, given that tendon extracellular matrix remodeling is altered with both age [25,67] and sex [25,72], ongoing studies are investigating altered regulatory pathways that drive divergent remodeling and functional outcomes in aged and female tendons.

The primary objective of this study was to apply step changes in tensile strain to define molecular and cellular programs by which healthy murine tendon explants adapt to increased (exercise) and decreased (disuse) strain magnitudes. Our findings reinforce that collagen incorporation and organization, not just synthesis, are critical determinants of tendon functional adaptation. We identify SLRPs, TGF-β isoforms, IL-6, and MMPs as strain-sensitive mediators that distinguish constructive, anabolic remodeling from degradative, catabolic responses. By linking these signaling molecules and matrix regulators to collagen structure and tissue mechanics in a controlled *ex vivo* model, we propose a mechanistic framework for load-dependent tendon adaptation (Figure 9). As tendon loading through physiotherapy remains one of the most effective treatments for tendinopathies [73], defining specific determinants of tendon adaptation has important therapeutic implications for optimizing rehabilitation protocols and identifying biologic therapies to enhance tendon regeneration. For instance, temporal modulation of TGF-β supplementation or early MMP inhibition combined with progressive loading may accelerate recovery of mechanical function more effectively than growth-factor delivery or loading alone. Importantly, characterizing these pathways in healthy tissues establishes foundation for developing targeted interventions to restore balanced remodeling in injured, diseased, or aged tendons. Together, this study provides a comprehensive, multiscale characterization of load-dependent tendon extracellular remodeling and highlights specific, targetable mediators that govern tendon adaptation.

## 4 Experimental Procedures

### 4.1 Explant Harvest and Culture

Flexor digitorum longus (FDL) tendon explants were harvested from 4-month (young) male C57BL/6J mice (Jackson Laboratories) under sterile conditions, as previously described [25]. Following extraction, explants were washed in 1x PBS supplemented with antibiotics and immediately transferred to culture medium. Explants were loaded in bioreactor grips at a 10 mm gauge length using a custom loading guide to ensure reproducibility [14]. To increase throughput, explants were “double-loaded,” with two tendons gripped together and cultured within a single well. Explants from the same mouse were not gripped together, and samples were randomized across culture wells. Explants were cultured in incubator-housed tensile loading bioreactors (Section 4.2) for up to two weeks at 37°C and 5% CO₂ under sterile conditions. The culture medium consisted of low glucose Dulbecco’s Modified Eagle Medium (DMEM; 1g/L glucose, Fisher Scientific) with 10% fetal bovine serum (Cytiva), 100 units/mL penicillin G, 100 μg/mL streptomycin (Fisher Scientific), and 0.25 μg/mL Amphotericin B (Sigma-Aldrich). Culture medium was replaced every two days.

### 4.2 *Ex Vivo* Tensile Loading

Tendon explants were subjected to *ex vivo* cyclic tensile strain using a custom-designed bioreactor system (Figure 1a) [14]. This custom-designed, incubator-housed bioreactor applied controlled displacements using a stepper motor linear actuator (Size 17-099; Haydon-Kerk) with tensile motion defined as positive (+) and compression motion defined as negative (-). Real-time load and displacement were measured using a precision load cell (5 lb capacity, Model 31; Honeywell) and a linear variable differential transformer (LVDT) (±0.1in, Model PLVX; Honeywell) with a sampling frequency of 10Hz. The cyclic tensile loading protocol consisted of a 20 g preload (defined as 0 µm) followed by a 1-hour displacement-controlled cyclic waveform (triangle or sinusoidal; 1 Hz) and subsequent 5-hour rest at slack (-100 µm relative displacement) (Supporting Figure S1). This 6-hour cycle was repeated four times per day throughout the culture period. The “zero displacement” point was redefined at each preload, and “zero load” was redefined at the end of each rest step. Stress-deprived tendons remained gripped but fully slack within the bioreactor system; therefore, real-time load and displacement data were not collected under stress-deprived conditions. All experiments experienced a final 1-hour 1% cyclic strain loading bout at the end of culture for mechanical characterization. Bioreactor control and waveform analysis were performed using custom MATLAB applications.

### 4.3 Quantitative Gene Expression

Tendon explants (n=10-15/group) were collected for qPCR analysis immediately following the final 1-hour loading bout at 1% cyclic strain. Explants were cut from bioreactor grips, rinsed in 1x PBS, flash-frozen in liquid nitrogen, and homogenized in TRIzol™ Reagent (Invitrogen) using a bead homogenizer (Benchmark Scientific). Homogenates were stored at −80°C until RNA extraction and purification was performed [26]. Total RNA was reverse transcribed to cDNA, and qualitative real-time polymerase chain reaction (PCR) was performed using the Bio-Rad CFX Opus 384 Real-Time PCR System with SYBR Green Power Up Master Mix (Applied Biosystems).

Gene expression was assessed for ECM proteins (*Col1a1, Col3a1, Dcn, Bgn, Fmod*), matrix degrading enzymes (*Mmp1, Mmp3, Mmp13*), secreted factors (*Il6, Tgfb1, Tnfa*), and apoptosis (*Casp3*). Murine primer sequences are listed in the Supporting Information (Supporting Table S1). Expression levels were calculated from the threshold cycle (*C*_t_) values and normalized to housekeeping gene *Actb* and to the day 7 average value using the 2^−ΔΔCt^ method [27]. Data are presented as log₂-transformed fold changes.

### 4.4 Matrix Biosynthesis and Tissue Composition

Synthesis of sulfated glycosaminoglycans (sGAG) and total protein (indicative of collagen synthesis) were measured by 24-hour incorporation of ^35^S-sulfate (2 μCi/mL) and ^3^H-proline (1 μCi/mL), respectively. Following radiolabel incorporation, tendons (n=10-16/group) were hydrated and weighed to obtain wet weight, then lyophilized for 3 hours to determine dry weight. Water content of the tendon was calculated as the difference in wet weight and dry weight divided by the dry weight and multiplied by 100. Tendons were digested in Proteinase K (5 mg/mL; Sigma) for 18 hours and stored at −20 °C until further assays were performed. Radiolabel incorporation was quantified using a liquid scintillation counter (PerkinElmer).

Additional biochemical assays were performed to quantify tissue composition [25]. sGAG content was measured using the dimethylmethylene blue (DMMB) assay and double-stranded DNA was quantified using the PicoGreen® dye-binding assay. A 100 µL aliquot of each digest was hydrolyzed in 12 M HCl, dried, resuspended, and assayed for total collagen content using the hydroxyproline (OHP) assay [25]. All biosynthesis and composition data were normalized to tissue dry weight to account for variability in sample size.

### 4.5 Second Harmonic Generation Imaging

Second-harmonic generation (SHG) imaging was performed to assess collagen fiber organization. Tendons (n=8-14/group) were mounted in 1x PBS between a glass microscope slide and #1 cover glass. Samples were imaged on a Bruker Investigator two-photon microscope with 1040 nm excitation laser and 525 ± 70 nm bandpass detection filter for SHG detection. Z-series image stacks were acquired at 3μm steps over a ∼60 μm depth using a 16x water-immersion objective at 2x digital zoom. Individual optical sections were extracted, brightness was adjusted, and images were converted to 8-bit prior to fiber distribution analysis using FiberFit™, a free web application for quantifying the material symmetry of fibrous networks [28].

### 4.6 Multiplex Immunoassays

Protein secretion was quantified using custom multiplex immunoassays (Meso Scale Discovery) on spent culture medium. Media was collected from bioreactor wells prior to media changes on days 6, 8, and 12 and stored at -20C. Media was not collected on day 14 due to radiolabel addition for biosynthesis assays (Section 4.4). A custom multiplex ELISA was developed to measure matrix metalloproteinases MMP-3, MMP-9, and MMP-13 simultaneously. MMP-9 was detected using the U-PLEX Mouse MMP-9 (total) antibody set, while MMP-3 was detected using the R-PLEX Mouse MMP-3 (total) antibody set (Meso Scale Discovery). The MMP-13 antibody set was purchased separately (Abcam; ab314458), conjugated in-house, and validated using a recombinant protein calibrator (Cusabio; CSB-EP014660MO; 20,000 pg/mL). The custom multiplex assay was performed per manufacture’s guidance using the UPLEX standard protocol. Substantial assay development and validation was performed to insure optimal assay performance and minimal antibody crosstalk. TIMP-1 was quantified separately using a single-plex immunoassay (R-PLEX Mouse TIMP-1 antibody set; Meso Scale Discovery). TGF- β family proteins (TGF-β1, TGF-β2, TGF-β3) were measured using U-PLEX Custom Biomarker Group 2 (mouse) Assay (Meso Scale Discovery) performed on acid-dissociated samples. Concentration values (n=5/group/timepoint for MMP/TIMP, n=9/group/timepoint for TGF-βs) represent the total protein concentration produced by two “double-loaded” tendon explants in a single bioreactor well. Concentrations are paired values as spent media was collected longitudinally over the culture period for each experiment. Data are presented as the log-transformed change in concentration post-intervention relative to pre-intervention for each sample.

### 4.7 MMP Activity

Generic MMP activity (MMPs 1, 2, 3, 7, 8, 9, 10, 13, and 14) was determined using a commercially available FRET-based generic MMP activity kit (SensoLyte 520 Generic MMP Activity Kit Fluorimetric; Anaspec) on spent culture medium (n = 4-8/group/time). MMP activity (MMPa) is represented as the concentration of MMP cleaved product (5-FAM-Pro-Leu-OH), the final product of the MMP enzymatic reaction.

### 4.8 Dynamic Mechanical Testing

Tendons analyzed for mechanical function were gripped with sandpaper at a 10-mm gauge length and loaded into a custom PBS-filled chamber integrated with a tensile testing frame (Instron) [29–31]. Tendons (n=18-23/group) were preloaded to 0.02 N and preconditioned with 10 cycles between 0.02 and 0.04 N. Following a 5-minute hold, tendons underwent three stress relaxations at 4, 6, and 8% grip strain at 5% s^−1^, with sinusoidal frequency sweeps superimposed on the static strain. Each frequency sweep consisted of 10 cycles of 0.125% amplitude sinusoidal grip strain at 0.1 and 1 Hz. After the relaxations, tendons were returned to zero displacement and subjected to a final ramp to failure at 0.1% per second. Load measurements were obtained using a 10 N load cell with 0.01 N resolution. Cross-sectional area was determined manually from post-preload images, assuming an elliptical tendon cross-section. Quasi-static and dynamic mechanical parameters were calculated using custom MATLAB software. Maximum stress (σ-max) and strain (ε-max) were determined from the peak of the stress–strain curve at sample failure. Stiffness (k) and elastic modulus (E) were calculated as the slope of the manually defined linear region of the load–displacement and stress–strain curves, respectively. Tissue stress relaxation (SR) was calculated as the percent change in stress under constant 4%, 6%, or 8% strain. Dynamic modulus (E*) and phase angle (δ) between stress and strain were extracted from sinusoidal frequency sweeps at each strain, as described previously [29,31,32]. Instron calibration for phase angle calculations was performed by subjecting a reference spring to dynamic frequency sweeps. A time delay correction factor was calculated at each frequency and applied to the time domain data so that the calibrated spring showed no measurable phase lag, consistent with purely elastic behavior.

### 4.9 Study Exclusion and Statistical Evaluation

A total of 25 bioreactor experiments were performed. All tendons within each bioreactor were subjected to a single strain profile, such that each bioreactor experiment represented a one loading intervention (control, disuse, or exercise), with samples (max 20/experiment) distributed across assays. Day 7 data was collected from a separate set of bioreactor experiments in which tendons were cultured at 1% cyclic strain for 7 days; therefore, the day 7 remodeling measurements are not paired with the day 14 data but serve as a reference for baseline tissue behavior prior to the intervention. Additional samples were collected at day 0 for comparison with *in vivo* native phenotypes (Supporting Figures S3 and S4).

Data sets were excluded from final analysis due to contamination or erroneous bioreactor loading. Erroneous loading events were identified though MATLAB analysis of real-time displacement and load data. Studies were excluded due to both under and overloading events that were attributed to internal (software, hardware) and external (environmental) factors. Data from a total of 17 studies is included in this manuscript (4-5 studies per loading condition). Even between successful experiments, we found high variability in measured loads (Figure 1d); therefore, data from multiple experiments were pooled for each assay when possible.

All data are presented as mean ± 95% confidence interval. Data points more than two standard deviations from the mean were removed as outliers. Statistical evaluation between loading conditions at day 14 was performed using one-way analysis of variance (ANOVA) and Bonferroni corrected post hoc *t*-tests versus control (GraphPad Prism). Additional one-tailed, t-tests were performed to identify significant adaptations between day 7 and day 14 for each loading intervention. Paired statistical tests were performed on media protein concentrations as samples were collected longitudinally throughout an experiment. For all comparisons, significance was noted at \**p* < 0.05 and a trend set at *p* < 0.10.

### 4.10 Correlation Analysis

A post-hoc correlation analysis was performed on 32 measured variables (Supporting Table S2). Correlations between variables could only be performed only when both measures were available for the same sample. For example, SHG imaging was performed prior to mechanical testing on a subset of explants, and spent culture medium analyzed for protein release was collected from wells corresponding to the tendons used for mechanical, biosynthesis/composition, or gene expression assays. Because of experimental constraints, not all parameters could be measured on every tendon. After outlier analysis (Section 4.10), data were pooled across loading interventions to generate a master data matrix for correlation. Of note, measured protein concentrations in medium were divided by two to estimate per-tendon release for double-loaded bioreactor wells (Section 4.7). A Pearson r correlation analysis was performed in GraphPad Prism. The full correlation matrix Pearson r values, p-values, and sample sizes are available in Supporting Information (Tables S3-S5). Simple linear regressions were also performed for paired variables of interest using GraphPad Prism. R^2^ goodness of fit and p-values for slope significance are shown with regression trendlines and 95% confidence intervals.

## 5 Acknowledgements

This study was supported by the National Institute of Health (K99/R00-AG063896, R35-GM151127), Wu Tsai Human Performance Alliance (2023 Agility Project Grant), Hevolution and the American Federation for Aging Research (2024 New Investigator Award in Aging Biology and Gerontology Research), National Science Foundation (GRFP; Stowe), and Neurophotonics Center at Boston University. The authors alone are responsible for the content and writing of the article. The authors would like to acknowledge Sam Mlawer for his assistance with bioreactor control and analysis software.

## 6 Conflict of Interest

The authors declare that the research was conducted in the absence of any commercial or financial relationships that could be construed as a potential conflict of interest.

## 7 Authors’ Contributions

EJ Stowe has contributed to all aspects of this study, including research design, data acquisition, interpretation/analysis of data, and drafting/revision of manuscript. BK Connizzo has contributed significantly to research design, interpretation/analysis of data, and drafting/revision of the manuscript. All authors have read and approved the final submitted manuscript.

## Notes

### Competing Interest Statement

The authors have declared no competing interest.

