## Supporting Information for "Regulators of ECM Structure Enable Functional Adaptation to Tensile Loading in Tendon Explants"

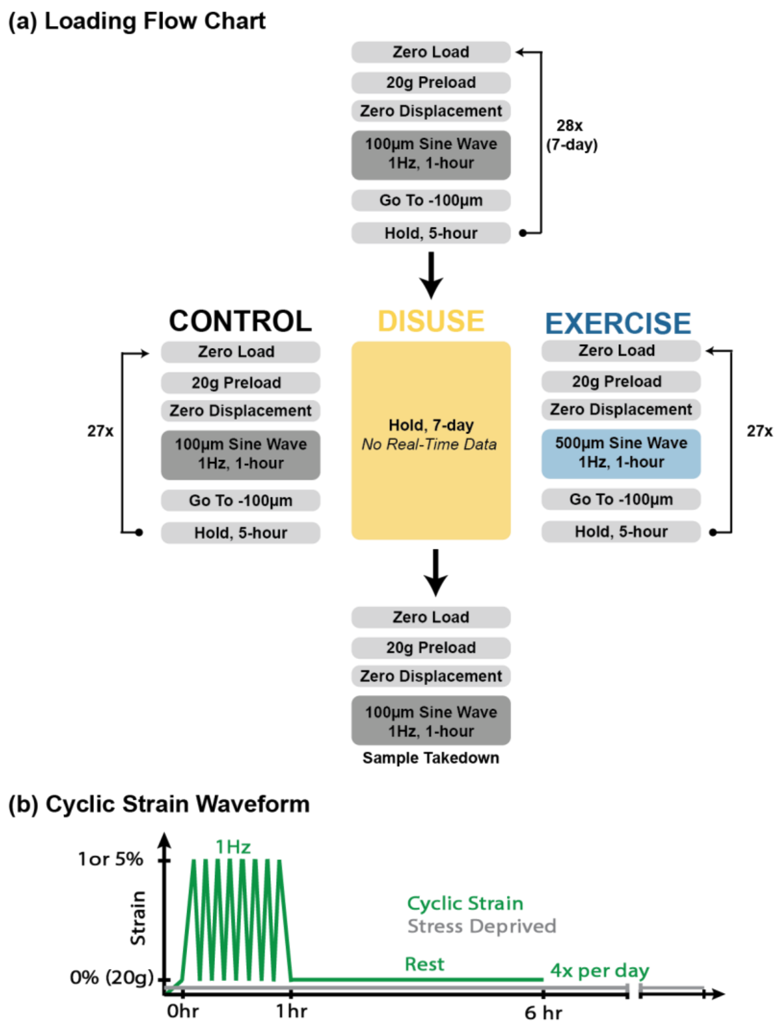


**Supporting Figure S1.** (a) Bioreactor loading flow chart and (b) cyclic waveform schematic.

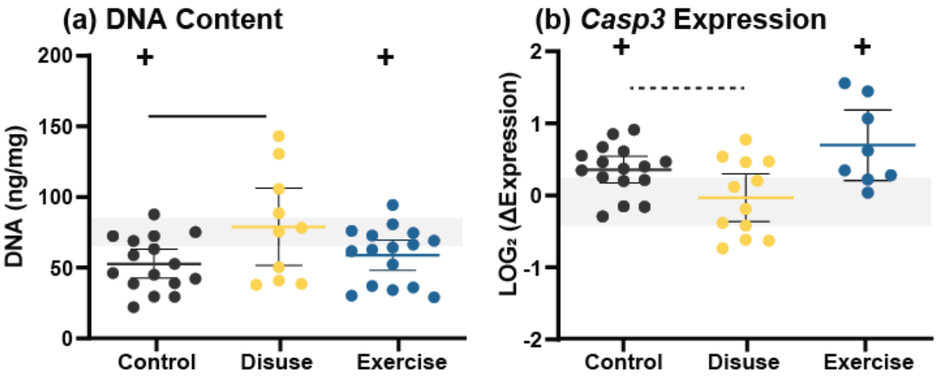


**Supporting Figure S2.** (a) DNA content and (b) *Casp3* gene expression of day 14 show no significant injury response for control, disuse, or exercise tendons. Gene expression data presented as relative expression to day 7 pre-intervention level. Data are shown as individual points with mean ± 95% confidence interval. Significant ($p < 0.05$) comparisons versus control are indicated with a solid line while trends ($p < 0.1$) are indicated with a dashed line. Grey horizontal bands show the day 7 pre-intervention reference levels (± 95% CI), and (+) symbols indicate significant differences from day 7 ($p < 0.05$).

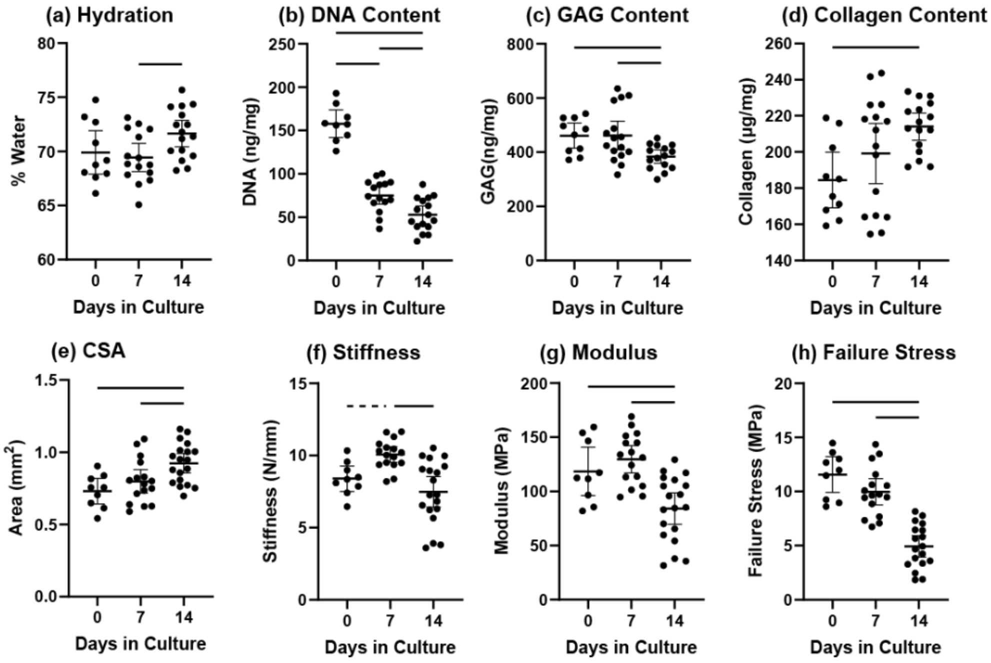


**Supporting Figure S3.** (a) Hydration, (b) DNA content, (c) GAG content, (d) collagen content, (e) CSA, (f) stiffness, (g) elastic modulus, and (h) failure stress of tendons at day 0 (native), day 7, and day 14 of *ex vivo* control loading (1% cyclic strain)**.** Data are shown as individual points with mean ± 95% confidence interval. Significant ($p < 0.05$) comparisons are indicated with a solid line while trends ($p < 0.1$) are indicated with a dashed line.

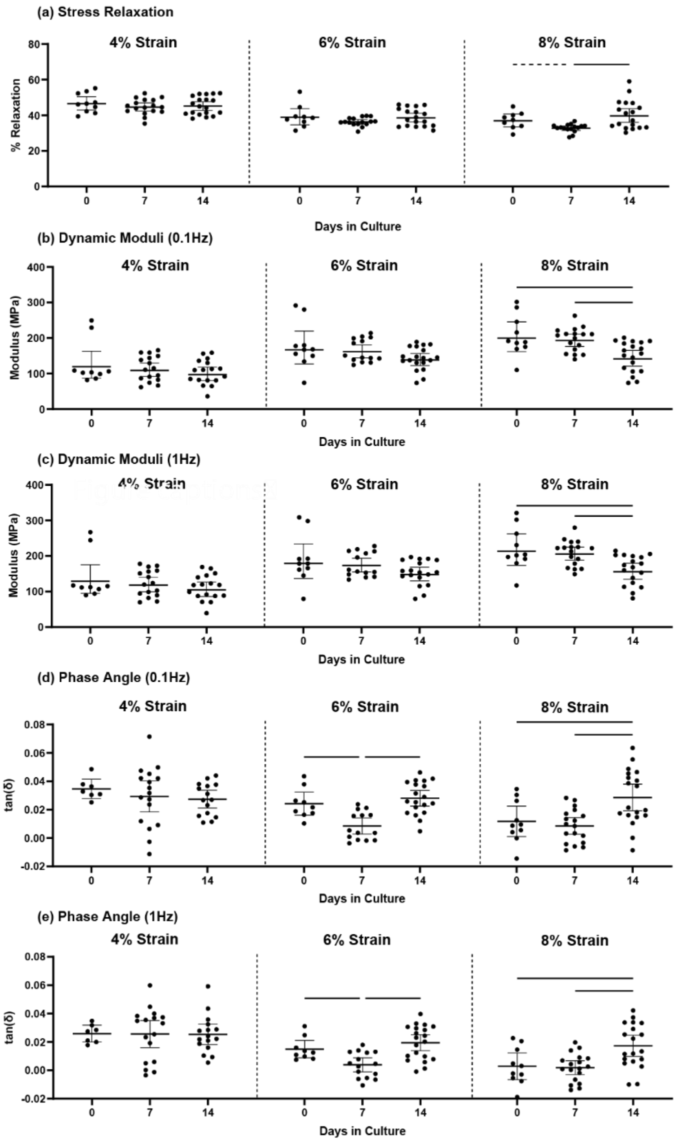


**Supporting Figure S4.** (a) Stress relaxation, (b,c) dynamic modulus, and (d,e) phase angle of tendons at day 0 (native), day 7, and day 14 of *ex vivo* control loading (1% cyclic strain)**.** Dynamic testing was conducted at 4%, 6%, and 8% grip strain with 0.1 Hz and 1Hz frequency sweeps. Data are shown as individual points with mean ± 95% confidence interval. Significant ($p < 0.05$) comparisons are indicated with a solid line while trends ($p < 0.1$) are indicated with a dashed line.

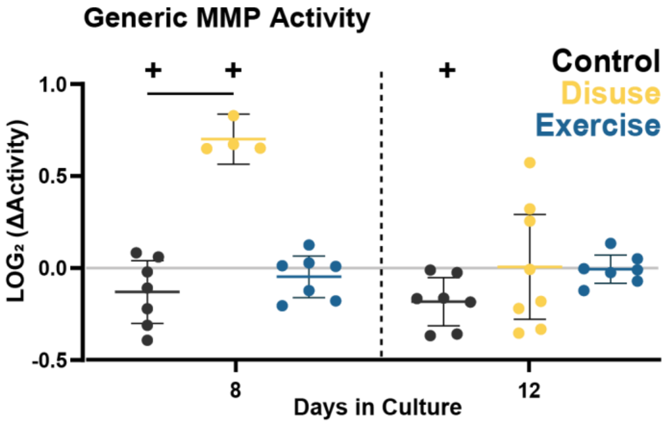


**Supporting Figure S5.** Fold change in generic MMP activity in spent culture medium at days 8 and 12 relative to day 6 pre-intervention levels. Activity measures are paired values collected longitudinally over the culture period for each experiment. Data are shown as individual points with mean ± 95% confidence interval. Significant ($p < 0.05$) comparisons versus control are indicated with a solid line. (+) symbols indicate significant differences from day 6 ($p < 0.05$). Raw data in Supporting Figure S6.

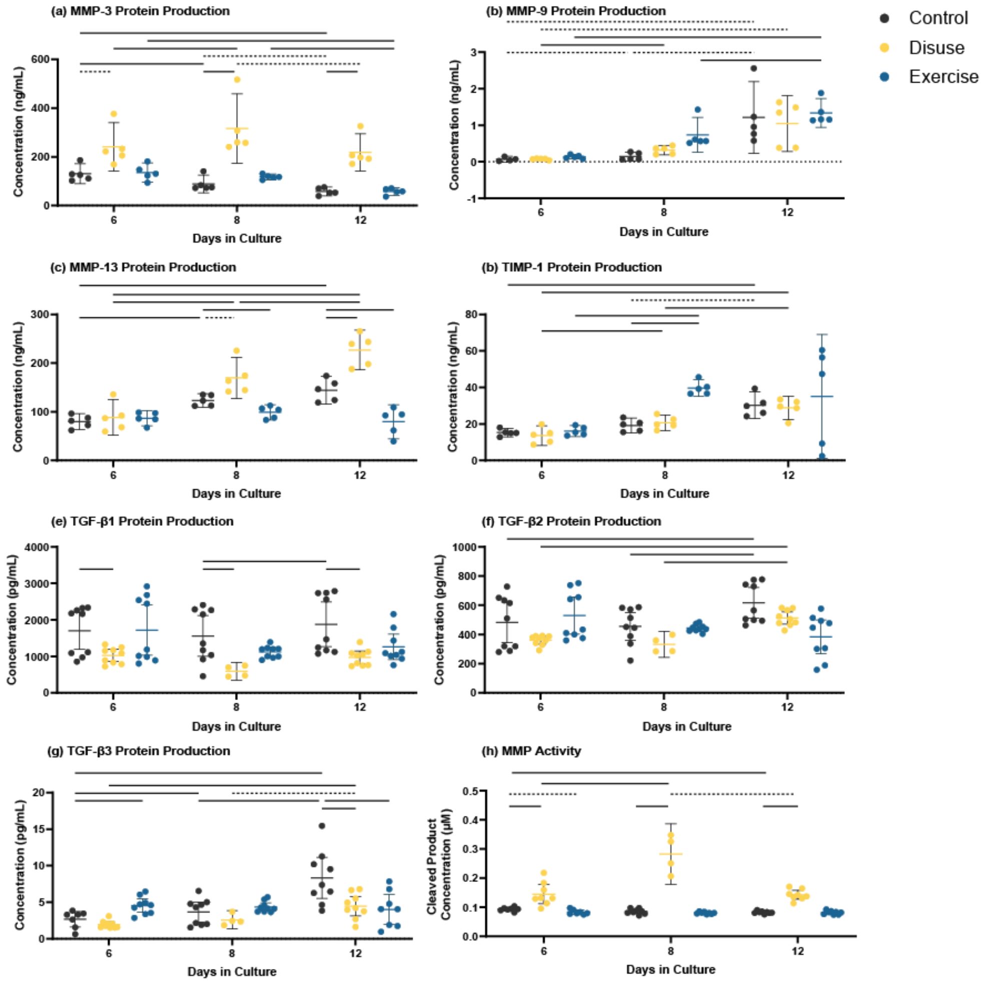


**Supporting Figure S6.** Concentrations of (a) MMP-3, (b) MMP-9, (c) MMP-13, (d) TIMP-1, (e) TGF-β1, (f) TGF-β2, and (g) TGF-β3 in spent culture medium at days 6, 8, and 12. (h) Concentration of cleaved MMP product representing MMP activity in culture medium. Concentrations are paired values collected longitudinally over the culture period for each experiment. Data are shown as individual points with mean ± 95% confidence interval. Multiple comparisons are performed with respect to both time and intervention. Significant ($p < 0.05$) pairwise comparisons are indicated with a solid line while trends ($p < 0.1$) are indicated with a dashed line.

**Supporting Table S1.** Murine gene names and primer forward/reverse sequences used for quantitative gene expression analysis.

| Target | Gene Name | Forward (5'-> 3') | Reverse (5'-> 3') |
|---|---|---|---|
| Beta Actin | *Actb*[1] | GGCTGTATTCCCCTCCATCG | CCAGTTGGTAACAATGCCATGT |
| Caspase-3 | *Casp3*[2] | ATGGGAGCAAGTCAGTGGAC | CGTACCAGAGCGAGATGACA |
| IL-6 | *Il6* [3] | TAGCTACCTGGAGTACATGAAGAACA | TGGTCCTTAGCCACTCCTTCTG |
| TNF-α | *Tnfa*[4] | GTCTCAGCCTCTTCTCATTC | CATAGAACTGATGAGAGGGA |
| Collagen 1 | *Col1a1*[3] | GACATGTTCAGCTTTGTGGACCTC | GGGACCCTTAGGCCATTGTGTA |
| Collagen 3 | *Col3a1*[5] | CAGAGCAACGGTCATACTCATTCACC | CAGCAACAGCAGAAGAGAAGCACC |
| Biglycan | *Bgn*[5] | TTTCTGAGCTTCGCAAGGATG | GGGCGTAGAGGTGCTGGAG |
| Decorin | *Dcn*[3] | CTATGTGCCCCTACCGATGC | CAGAACACTGCACCACTCGAAG |
| Fibromodulin | *Fmod*[3] | CTCCAACCCAAGGAGACCAG | GGATCCACCAGTGAGAGTCTTC |
| TGF-β1 | *Tgfb1*[6] | ATTCCTGGCGTTACCTTGG | CCTGTATTCCGTCTCCTTGG |
| MMP-1 | *Mmp1*[7] | TCAACCAGGCCAAGGTATTG | ATGAGCAGCCACGAGAAATAG |
| MMP-3 | *Mmp3*[2] | ACATGGAGACTTTGTCCCTTTTG | TTGGCTGAGTGGTAGAGTCCC |
| MMP-13 | *Mmp13*[2] | TCAGTCTCTTCACCTCTTTTGGGAATCC | TCAGTTTCTTTATGGTCCAGGCGATG |

**Supporting Table S2:** Details of variables included in the correlation analysis.

| Variable | Full Name | Units | Assay | Timepoint |
|---|---|---|---|---|
| E | Elastic Modulus | Mpa | Mechanical Testing | Day 14 |
| σ-max | Maximum Stress | MPa | Mechanical Testing | Day 14 |
| ε-max | Maximum Strain | % | Mechanical Testing | Day 14 |
| E* | Dynamic Modulus | MPa | Mechanical Testing | Day 14 |
| SR | Stress Relaxation | % | Mechanical Testing | Day 14 |
| Tan δ | Tangent of Phase Angle | - | Mechanical Testing | Day 14 |
| AI | Collagen Alignment Index | [0,1] | SHG Imaging | Day 14 |
| $^{3}H$ | 3H-Proline Incorporation | pmol/hr/mg | Biosynthesis/Composition | Day 14 |
| COL | Collagen Content | µg/mg | Biosynthesis/Composition | Day 14 |
| $^{35}S$ | 35S-Sulfate Incorporation | pmol/hr/mg | Biosynthesis/Composition | Day 14 |
| GAG | Glycosaminoglycan Content | ng/mg | Biosynthesis/Composition | Day 14 |
| $H_20$ | Water Content | % | Biosynthesis/Composition | Day 14 |
| [TGF-β1] | TGF-β1 Concentration in Media | pg/mL | Multiplex Immunoassay | Day 12 |
| [TGF-β2] | TGF-β2 Concentration in Media | pg/mL | Multiplex Immunoassay | Day 12 |
| [TGF-β3] | TGF-β3 Concentration in Media | pg/mL | Multiplex Immunoassay | Day 12 |
| [MMP-3] | MMP-3 Concentration in Media | ng/mL | Multiplex Immunoassay | Day 12 |
| [MMP-9] | MMP-9 Concentration in Media | ng/mL | Multiplex Immunoassay | Day 12 |
| [MMP-13] | MMP-13 Concentration in Media | ng/mL | Multiplex Immunoassay | Day 12 |
| [TIMP-1] | TIMP-1 Concentration in Media | ng/mL | Immunoassay | Day 12 |
| MMPa | Generic MMP Activity in Media | Cleavage Product µM | Activity Assay | Day 12 |
| *Col1* | *Col1* Gene Expression | Relative Expression (*Actb*) | qPCR | Day 14 |
| *Col3* | *Col3* Gene Expression | Relative Expression (*Actb*) | qPCR | Day 14 |
| *Bgn* | *Bgn* Gene Expression | Relative Expression (*Actb*) | qPCR | Day 14 |
| *Fmod* | *Fmod* Gene Expression | Relative Expression (*Actb*) | qPCR | Day 14 |
| *Dcn* | *Dcn* Gene Expression | Relative Expression (*Actb*) | qPCR | Day 14 |
| *Mmp1* | *Mmp1* Gene Expression | Relative Expression (*Actb*) | qPCR | Day 14 |
| *Mmp3* | *Mmp3* Gene Expression | Relative Expression (*Actb*) | qPCR | Day 14 |
| *Mmp13* | *Mmp13* Gene Expression | Relative Expression (*Actb*) | qPCR | Day 14 |
| *Tgfb1* | *Tgfb1* Gene Expression | Relative Expression (*Actb*) | qPCR | Day 14 |
| *Il6* | *Il6* Gene Expression | Relative Expression (*Actb*) | qPCR | Day 14 |
| *Tnfa1* | *Tnfa1* Gene Expression | Relative Expression (*Actb*) | qPCR | Day 14 |
| *Casp3* | *Casp3* Gene Expression | Relative Expression (*Actb*) | qPCR | Day 14 |

**Supporting Table S3.** Pearson r correlation matrix.

| Pearson r | E | σ-max | ε-max | E* | SR | Tan δ | AI | $^{3}H$ | COL | $^{35}S$ | GAG | $H_20$ | [TGF-β1] | [TGF-β2] | [TGF-β3] | [MMP-3] | [MMP-9] | [MMP-13] | [TIMP-1] | *Col1* | *Col3* | *Bgn* | *Fmod* | *Dcn* | *Mmp1* | *Mmp3* | *Mmp13* | *Tgfb1* | *Il6* | *Tnfa1* | *Casp3* |
|---|---|---|---|---|---|---|---|---|---|---|---|---|---|---|---|---|---|---|---|---|---|---|---|---|---|---|---|---|---|---|---|
| E | 1.0000 | 0.9001 | 0.2889 | 0.7655 | -0.5997 | -0.0080 | 0.5427 | | | | | | -0.4082 | -0.4366 | -0.5303 | 0.0037 | -0.1183 | -0.3580 | -0.5373 | | | | | | | | | | | | |
| σ-max | 0.9001 | 1.0000 | 0.4979 | 0.6580 | -0.5994 | 0.0317 | 0.6165 | | | | | | -0.4355 | -0.5060 | -0.6003 | 0.0013 | -0.2078 | -0.3320 | -0.5338 | | | | | | | | | | | | |
| ε-max | 0.2889 | 0.4979 | 1.0000 | -0.1273 | -0.0813 | 0.0798 | 0.6190 | | | | | | -0.2995 | -0.4757 | -0.4853 | -0.1254 | -0.5039 | -0.3564 | -0.5502 | | | | | | | | | | | | |
| E* | 0.7655 | 0.6580 | -0.1273 | 1.0000 | -0.5695 | -0.0609 | 0.3450 | | | | | | -0.4715 | -0.4361 | -0.5299 | 0.0943 | 0.0825 | -0.2388 | -0.3391 | | | | | | | | | | | | |
| SR | -0.5997 | -0.5994 | -0.0813 | -0.5695 | 1.0000 | 0.1578 | -0.4924 | | | | | | 0.6056 | 0.8932 | 0.6675 | 0.5883 | 0.3132 | 0.7846 | 0.8171 | | | | | | | | | | | | |
| Tan δ | -0.0080 | 0.0317 | 0.0798 | -0.0609 | 0.1578 | 1.0000 | -0.5078 | | | | | | -0.0762 | 0.1236 | 0.0626 | 0.2383 | -0.0546 | 0.4379 | 0.3894 | | | | | | | | | | | | |
| AI | 0.5427 | 0.6165 | 0.6190 | 0.3450 | -0.4924 | -0.5078 | 1.0000 | | | | | | -0.3131 | -0.6284 | -0.7530 | -0.7511 | -0.2028 | -0.9246 | -0.8852 | | | | | | | | | | | | |
| $^{3}H$ | | | | | | | | 1.0000 | 0.4979 | 0.5456 | -0.0872 | 0.0888 | 0.1988 | -0.4375 | -0.2432 | -0.4670 | 0.4187 | -0.7347 | 0.9058 | | | | | | | | | | | | |
| COL | | | | | | | | 0.4979 | 1.0000 | 0.1736 | -0.3503 | 0.0035 | 0.2213 | -0.0673 | 0.2666 | -0.3526 | 0.2177 | -0.5285 | 0.7198 | | | | | | | | | | | | |
| $^{35}S$ | | | | | | | | 0.5456 | 0.1736 | 1.0000 | 0.2299 | 0.0827 | 0.1662 | -0.0797 | -0.2665 | -0.4700 | 0.4348 | -0.5081 | 0.6140 | | | | | | | | | | | | |
| GAG | | | | | | | | -0.0872 | -0.3503 | 0.2299 | 1.0000 | 0.2487 | -0.6122 | 0.2980 | -0.1424 | 0.7491 | 0.2126 | 0.6361 | -0.2345 | | | | | | | | | | | | |
| $H_20$ | | | | | | | | 0.0888 | 0.0035 | 0.0827 | 0.2487 | 1.0000 | -0.4120 | -0.0473 | -0.3102 | 0.6686 | 0.2836 | 0.4271 | 0.0056 | | | | | | | | | | | | |
| [TGF-β1] | -0.4082 | -0.4355 | -0.2995 | -0.4715 | 0.6056 | -0.0762 | -0.3131 | 0.1988 | 0.2213 | 0.1662 | -0.6122 | -0.4120 | 1.0000 | 0.7239 | 0.7131 | -0.1591 | 0.0783 | -0.0269 | 0.2610 | 0.6675 | 0.3640 | 0.6269 | 0.6102 | 0.0702 | 0.5489 | -0.6856 | -0.5699 | 0.7844 | -0.3731 | 0.3403 | -0.0950 |
| [TGF-β2] | -0.4366 | -0.5060 | -0.4757 | -0.4361 | 0.8932 | 0.1236 | -0.6284 | -0.4375 | -0.0673 | -0.0797 | 0.2980 | -0.0473 | 0.7239 | 1.0000 | 0.7858 | 0.4285 | -0.2073 | 0.7541 | 0.5264 | 0.4002 | -0.0875 | 0.7310 | 0.6532 | -0.2359 | 0.8674 | -0.6121 | -0.3576 | 0.8175 | -0.3701 | 0.5331 | 0.1663 |
| [TGF-β3] | -0.5303 | -0.6003 | -0.4853 | -0.5299 | 0.6675 | 0.0626 | -0.7530 | -0.2432 | 0.2666 | -0.2665 | -0.1424 | -0.3102 | 0.7131 | 0.7858 | 1.0000 | -0.2397 | -0.2375 | 0.1377 | 0.0591 | 0.6289 | 0.0358 | 0.7098 | 0.7044 | -0.0262 | 0.8951 | -0.5902 | -0.4731 | 0.8400 | -0.3989 | 0.3972 | 0.2891 |
| [MMP-3] | 0.0037 | 0.0013 | -0.1254 | 0.0943 | 0.5883 | 0.2383 | -0.7511 | -0.4670 | -0.3526 | -0.4700 | 0.7491 | 0.6686 | -0.1591 | 0.4285 | -0.2397 | 1.0000 | -0.0922 | 0.8559 | -0.0332 | | | | | | | | | | | | |
| [MMP-9] | -0.1183 | -0.2078 | -0.5039 | 0.0825 | 0.3132 | -0.0546 | -0.2028 | 0.4187 | 0.2177 | 0.4348 | 0.2126 | 0.2836 | 0.0783 | -0.2073 | -0.2375 | -0.0922 | 1.0000 | -0.1282 | -0.1996 | | | | | | | | | | | | |
| [MMP-13] | -0.3580 | -0.3320 | -0.3564 | -0.2388 | 0.7846 | 0.4379 | -0.9246 | -0.7347 | -0.5285 | -0.5081 | 0.6361 | 0.4271 | -0.0269 | 0.7541 | 0.1377 | 0.8559 | -0.1282 | 1.0000 | 0.0464 | | | | | | | | | | | | |
| [TIMP-1] | -0.5373 | -0.5338 | -0.5502 | -0.3391 | 0.8171 | 0.3894 | -0.8852 | 0.9058 | 0.7198 | 0.6140 | -0.2345 | 0.0056 | 0.2610 | 0.5264 | 0.0591 | -0.0332 | -0.1996 | 0.0464 | 1.0000 | | | | | | | | | | | | |
| *Col1* | | | | | | | | | | | | | 0.6675 | 0.4002 | 0.6289 | | | | | 1.0000 | 0.6658 | 0.4633 | 0.6255 | -0.2647 | -0.1692 | -0.5030 | -0.4503 | -0.0489 | -0.4313 | 0.1036 | -0.1874 |
| *Col3* | | | | | | | | | | | | | 0.3640 | -0.0875 | 0.0358 | | | | | 0.6658 | 1.0000 | 0.1077 | 0.2590 | -0.4754 | -0.4647 | -0.6720 | -0.4333 | -0.4229 | -0.6381 | -0.0240 | -0.4293 |
| *Bgn* | | | | | | | | | | | | | 0.6269 | 0.7310 | 0.7098 | | | | | 0.4633 | 0.1077 | 1.0000 | 0.7581 | 0.2626 | 0.0297 | 0.1588 | 0.0779 | 0.6296 | 0.2009 | -0.1741 | 0.3818 |
| *Fmod* | | | | | | | | | | | | | 0.6102 | 0.6532 | 0.7044 | | | | | 0.6255 | 0.2590 | 0.7581 | 1.0000 | 0.1679 | -0.2038 | -0.0404 | -0.0820 | 0.5611 | 0.0200 | -0.1467 | 0.2980 |
| *Dcn* | | | | | | | | | | | | | 0.0702 | -0.2359 | -0.0262 | | | | | -0.2647 | -0.4754 | 0.2626 | 0.1679 | 1.0000 | 0.6329 | 0.9063 | 0.5688 | 0.7563 | 0.9102 | -0.4291 | 0.7348 |
| *Mmp1* | | | | | | | | | | | | | 0.5489 | 0.8674 | 0.8951 | | | | | -0.1692 | -0.4647 | 0.0297 | -0.2038 | 0.6329 | 1.0000 | 0.5365 | 0.4633 | 0.1960 | 0.5074 | -0.0956 | 0.2568 |
| *Mmp3* | | | | | | | | | | | | | -0.6856 | -0.6121 | -0.5902 | | | | | -0.5030 | -0.6720 | 0.1588 | -0.0404 | 0.9063 | 0.5365 | 1.0000 | 0.6529 | 0.7178 | 0.9710 | -0.4236 | 0.7395 |
| *Mmp13* | | | | | | | | | | | | | -0.5699 | -0.3576 | -0.4731 | | | | | -0.4503 | -0.4333 | 0.0779 | -0.0820 | 0.5688 | 0.4633 | 0.6529 | 1.0000 | 0.3113 | 0.5503 | -0.1836 | 0.3864 |
| *Tgfb1* | | | | | | | | | | | | | 0.7844 | 0.8175 | 0.8400 | | | | | -0.0489 | -0.4229 | 0.6296 | 0.5611 | 0.7563 | 0.1960 | 0.7178 | 0.3113 | 1.0000 | 0.7831 | -0.3401 | 0.7297 |
| *Il6* | | | | | | | | | | | | | -0.3731 | -0.3701 | -0.3989 | | | | | -0.4313 | -0.6381 | 0.2009 | 0.0200 | 0.9102 | 0.5074 | 0.9710 | 0.5503 | 0.7831 | 1.0000 | -0.4021 | 0.6855 |
| *Tnfa1* | | | | | | | | | | | | | 0.3403 | 0.5331 | 0.3972 | | | | | 0.1036 | -0.0240 | -0.1741 | -0.1467 | -0.4291 | -0.0956 | -0.4236 | -0.1836 | -0.3401 | -0.4021 | 1.0000 | -0.3328 |
| *Casp3* | | | | | | | | | | | | | -0.0950 | 0.1663 | 0.2891 | | | | | -0.1874 | -0.4293 | 0.3818 | 0.2980 | 0.7348 | 0.2568 | 0.7395 | 0.3864 | 0.7297 | 0.6855 | -0.3328 | 1.0000 |

**Supporting Table S4.** Correlation analysis p-values.

| p-value | E | σ-max | ε-max | E* | SR | Tan δ | AI | $^3$H | COL | $^{35}$S | GAG | $H_2$0 | [TGF-β1] | [TGF-β2] | [TGF-β3] | [MMP-3] | [MMP-9] | [MMP-13] | [TIMP-1] | *Col1* | *Col3* | *Bgn* | *Fmod* | *Dcn* | *Mmp1* | *Mmp3* | *Mmp13* | *Tgfb1* | *Il6* | *Tnfa1* | *Casp3* |
|---|---|---|---|---|---|---|---|---|---|---|---|---|---|---|---|---|---|---|---|---|---|---|---|---|---|---|---|---|---|---|---|
| E | | 1.1E-19 | 0.041839 | 2E-10 | 4.2E-06 | 0.957058 | 0.02 | | | | | | 0.09262 | 0.0700963 | 0.0346017 | 0.9908556 | 0.7141658 | 0.25327253 | 0.0716167 | | | | | | | | | | | | |
| σ-max | 1E-19 | | 0.000273 | 5E-07 | 5.4E-06 | 0.832537 | 0.0064 | | | | | | 0.070885 | 0.0321389 | 0.0139378 | 0.9968796 | 0.5169079 | 0.29170462 | 0.0738287 | | | | | | | | | | | | |
| ε-max | 0.0418 | 0.00027 | | 0.3994 | 0.58299 | 0.597948 | 0.0062 | | | | | | 0.242843 | 0.0535806 | 0.0667058 | 0.7133539 | 0.1140506 | 0.28206583 | 0.0794557 | | | | | | | | | | | | |
| E* | 2E-10 | 5E-07 | 0.399363 | | 2.9E-05 | 0.680743 | 0.1906 | | | | | | 0.065258 | 0.0913203 | 0.0512924 | 0.7826908 | 0.8095466 | 0.47954137 | 0.3076544 | | | | | | | | | | | | |
| SR | 4E-06 | 5.4E-06 | 0.582986 | 3E-05 | | 0.289595 | 0.0446 | | | | | | 0.009979 | 1.394E-06 | 0.0065546 | 0.0569281 | 0.348368 | 0.00423827 | 0.0021368 | | | | | | | | | | | | |
| Tan δ | 0.9571 | 0.83254 | 0.597948 | 0.6807 | 0.28959 | | 0.0375 | | | | | | 0.779124 | 0.6482943 | 0.8315348 | 0.5073042 | 0.8808293 | 0.20560672 | 0.266075 | | | | | | | | | | | | |
| AI | 0.02 | 0.00644 | 0.006159 | 0.1906 | 0.04465 | 0.037455 | | | | | | | 0.321683 | 0.0286308 | 0.0119282 | 0.0317093 | 0.6300142 | 0.00101115 | 0.0034676 | | | | | | | | | | | | |
| $^3$H | | | | | | | | | 0.002 | 0.000479 | 0.613223 | 0.60643 | 0.428994 | 0.0693909 | 0.3307877 | 0.0681751 | 0.1064985 | 0.00118774 | 1.368E-06 | | | | | | | | | | | | |
| COL | | | | | | | | 0.002 | | 0.311234 | 0.033526 | 0.983164 | 0.393223 | 0.7973555 | 0.3009257 | 0.1973534 | 0.4357642 | 0.04283645 | 0.0024815 | | | | | | | | | | | | |
| $^{35}$S | | | | | | | | 0.000479 | 0.3112 | | 0.177315 | 0.631558 | 0.509695 | 0.7532874 | 0.2850169 | 0.0661885 | 0.0923293 | 0.04450326 | 0.0113968 | | | | | | | | | | | | |
| GAG | | | | | | | | 0.613223 | 0.0335 | 0.177315 | | 0.132187 | 0.006928 | 0.2297773 | 0.57294 | 0.0008387 | 0.4291615 | 0.00808131 | 0.3819891 | | | | | | | | | | | | |
| $H_2$0 | | | | | | | | 0.60643 | 0.9832 | 0.631558 | 0.132187 | | 0.089325 | 0.8520696 | 0.2102412 | 0.0046317 | 0.2871768 | 0.09895669 | 0.9835437 | | | | | | | | | | | | |
| [TGF-β1] | 0.0926 | 0.07088 | 0.242843 | 0.0653 | 0.00998 | 0.779124 | 0.3217 | 0.428994 | 0.3932 | 0.509695 | 0.006928 | 0.089325 | | 9.046E-10 | 4.317E-09 | 0.4011348 | 0.6807445 | 0.8879046 | 0.1635467 | 0.009103 | 0.1657 | 0.00935 | 0.0157 | 0.79606 | 0.33801 | 0.0048 | 0.02117 | 0.000536 | 0.1708 | 0.2339 | 0.72624 |
| [TGF-β2] | 0.0701 | 0.03214 | 0.053581 | 0.0913 | 1.4E-06 | 0.648294 | 0.0286 | 0.069391 | 0.7974 | 0.753287 | 0.229777 | 0.85207 | 9.05E-10 | | 8.499E-12 | 0.0181476 | 0.2717502 | 1.4896E-06 | 0.0028046 | 0.156187 | 0.7474 | 0.0013 | 0.00828 | 0.37899 | 0.05681 | 0.0153 | 0.17387 | 0.000197 | 0.17455 | 0.0497 | 0.53831 |
| [TGF-β3] | 0.0346 | 0.01394 | 0.066706 | 0.0513 | 0.00655 | 0.831535 | 0.0119 | 0.330788 | 0.3009 | 0.285017 | 0.57294 | 0.210241 | 4.32E-09 | 8.499E-12 | | 0.2192197 | 0.2235413 | 0.48456129 | 0.764962 | 0.015985 | 0.8952 | 0.00207 | 0.00337 | 0.92322 | 0.04012 | 0.0206 | 0.06418 | 8.9E-05 | 0.14074 | 0.1596 | 0.27744 |
| [MMP-3] | 0.9909 | 0.99688 | 0.713354 | 0.7827 | 0.05693 | 0.507304 | 0.0317 | 0.068175 | 0.1974 | 0.066188 | 0.000839 | 0.004632 | 0.401135 | 0.0181476 | 0.2192197 | | 0.6279739 | 1.6529E-09 | 0.8617561 | | | | | | | | | | | | |
| [MMP-9] | 0.7142 | 0.51691 | 0.114051 | 0.8095 | 0.34837 | 0.880829 | 0.63 | 0.106499 | 0.4358 | 0.092329 | 0.429162 | 0.287177 | 0.680744 | 0.2717502 | 0.2235413 | 0.6279739 | | 0.4995964 | 0.2903664 | | | | | | | | | | | | |
| [MMP-13] | 0.2533 | 0.2917 | 0.282066 | 0.4795 | 0.00424 | 0.205607 | 0.001 | 0.001188 | 0.0428 | 0.044503 | 0.008081 | 0.098957 | 0.887905 | 1.49E-06 | 0.4845613 | 1.653E-09 | 0.4995964 | | 0.8076907 | | | | | | | | | | | | |
| [TIMP-1] | 0.0716 | 0.07383 | 0.079456 | 0.3077 | 0.00214 | 0.266075 | 0.0035 | 1.37E-06 | 0.0025 | 0.011397 | 0.381989 | 0.983544 | 0.163547 | 0.0028046 | 0.764962 | 0.8617561 | 0.2903664 | 0.80769067 | | | | | | | | | | | | | |
| *Col1* | | | | | | | | | | | | | 0.009103 | 0.1561872 | 0.0159851 | | | | | | 2E-05 | 0.00507 | 9.9E-05 | 0.13035 | 0.48866 | 0.0029 | 0.00753 | 0.780339 | 0.01221 | 0.5661 | 0.28855 |
| *Col3* | | | | | | | | | | | | | 0.165697 | 0.7473934 | 0.8951763 | | | | | 1.73E-05 | | 0.53183 | 0.13912 | 0.0039 | 0.04499 | 1E-05 | 0.00932 | 0.011375 | 4.9E-05 | 0.8926 | 0.01007 |
| *Bgn* | | | | | | | | | | | | | 0.009349 | 0.0012953 | 0.00207 | | | | | 0.005067 | 0.5318 | | 1.3E-07 | 0.12175 | 0.90389 | 0.3621 | 0.65148 | 3.89E-05 | 0.24725 | 0.3172 | 0.02157 |
| *Fmod* | | | | | | | | | | | | | 0.015695 | 0.0082759 | 0.003372 | | | | | 9.92E-05 | 0.1391 | 1.3E-07 | | 0.34252 | 0.40255 | 0.8235 | 0.64494 | 0.000556 | 0.91206 | 0.4154 | 0.08206 |
| *Dcn* | | | | | | | | | | | | | 0.796062 | 0.3789921 | 0.9232208 | | | | | 0.130347 | 0.0039 | 0.12175 | 0.34252 | | 0.00482 | 7E-14 | 0.0003 | 1.49E-07 | 8.5E-14 | 0.0113 | 5E-07 |
| *Mmp1* | | | | | | | | | | | | | 0.338006 | 0.0568061 | 0.0401184 | | | | | 0.488658 | 0.045 | 0.90389 | 0.40255 | 0.00482 | | 0.0217 | 0.05284 | 0.421267 | 0.02657 | 0.706 | 0.28848 |
| *Mmp3* | | | | | | | | | | | | | 0.004784 | 0.0152947 | 0.0205531 | | | | | 0.00285 | 1E-05 | 0.36207 | 0.82349 | 6.9E-14 | 0.02171 | | 2.1E-05 | 1.76E-06 | 1.9E-21 | 0.014 | 5.8E-07 |
| *Mmp13* | | | | | | | | | | | | | 0.021169 | 0.1738676 | 0.06418 | | | | | 0.007533 | 0.0093 | 0.65148 | 0.64494 | 0.0003 | 0.05284 | 2E-05 | | 0.068676 | 0.00075 | 0.2987 | 0.02187 |
| *Tgfb1* | | | | | | | | | | | | | 0.000536 | 0.0001973 | 8.896E-05 | | | | | 0.780339 | 0.0114 | 3.9E-05 | 0.00056 | 1.5E-07 | 0.42127 | 2E-06 | 0.06868 | | 4.4E-08 | 0.0491 | 6.6E-07 |
| *Il6* | | | | | | | | | | | | | 0.170805 | 0.1745516 | 0.1407373 | | | | | 0.012209 | 5E-05 | 0.24725 | 0.91206 | 8.5E-14 | 0.02657 | 2E-21 | 0.00075 | 4.4E-08 | | 0.0204 | 7.7E-06 |
| *Tnfa1* | | | | | | | | | | | | | 0.233857 | 0.0496583 | 0.1596318 | | | | | 0.566114 | 0.8926 | 0.31723 | 0.41536 | 0.01134 | 0.70605 | 0.014 | 0.2987 | 0.049091 | 0.02036 | | 0.05445 |
| *Casp3* | | | | | | | | | | | | | 0.726242 | 0.5383115 | 0.2774359 | | | | | 0.288554 | 0.0101 | 0.02157 | 0.08206 | 5E-07 | 0.28848 | 6E-07 | 0.02187 | 6.56E-07 | 7.7E-06 | 0.0544 | |

**Supporting Table S5.** Correlation analysis sample sizes (n).

| n | E | σ-max | ε-max | E* | SR | Tan δ | AI | $^3$H | COL | $^{35}$S | GAG | $H_20$ | [TGF-β1] | [TGF-β2] | [TGF-β3] | [MMP-3] | [MMP-9] | [MMP-13] | [TIMP-1] | *Col1* | *Col3* | *Bgn* | *Fmod* | *Dcn* | *Mmp1* | *Mmp3* | *Mmp13* | *Tgfb1* | *Il6* | *Tnfa1* | *Casp3* |
|---|---|---|---|---|---|---|---|---|---|---|---|---|---|---|---|---|---|---|---|---|---|---|---|---|---|---|---|---|---|---|---|
| E | 53 | 52 | 50 | 48 | 50 | 48 | 18 | | | | | | 18 | 18 | 16 | 12 | 12 | 12 | 12 | | | | | | | | | | | | |
| σ-max | 52 | 52 | 49 | 47 | 49 | 47 | 18 | | | | | | 18 | 18 | 16 | 12 | 12 | 12 | 12 | | | | | | | | | | | | |
| ε-max | 50 | 49 | 50 | 46 | 48 | 46 | 18 | | | | | | 17 | 17 | 15 | 11 | 11 | 11 | 11 | | | | | | | | | | | | |
| E* | 48 | 47 | 46 | 49 | 47 | 48 | 16 | | | | | | 16 | 16 | 14 | 11 | 11 | 11 | 11 | | | | | | | | | | | | |
| SR | 50 | 49 | 48 | 47 | 51 | 47 | 17 | | | | | | 17 | 17 | 15 | 11 | 11 | 11 | 11 | | | | | | | | | | | | |
| Tan δ | 48 | 47 | 46 | 48 | 47 | 51 | 17 | | | | | | 16 | 16 | 14 | 10 | 10 | 10 | 10 | | | | | | | | | | | | |
| AI | 18 | 18 | 18 | 16 | 17 | 17 | 36 | | | | | | 12 | 12 | 10 | 8 | 8 | 8 | 8 | | | | | | | | | | | | |
| $^3$H | | | | | | | | 37 | 36 | 37 | 36 | 36 | 18 | 18 | 18 | 16 | 16 | 16 | 16 | | | | | | | | | | | | |
| COL | | | | | | | | 36 | 40 | 36 | 37 | 39 | 17 | 17 | 17 | 15 | 15 | 15 | 15 | | | | | | | | | | | | |
| $^{35}$S | | | | | | | | 37 | 36 | 37 | 36 | 36 | 18 | 18 | 18 | 16 | 16 | 16 | 16 | | | | | | | | | | | | |
| GAG | | | | | | | | 36 | 37 | 36 | 39 | 38 | 18 | 18 | 18 | 16 | 16 | 16 | 16 | | | | | | | | | | | | |
| $H_20$ | | | | | | | | 36 | 39 | 36 | 38 | 41 | 18 | 18 | 18 | 16 | 16 | 16 | 16 | | | | | | | | | | | | |
| [TGF-β1] | 18 | 18 | 17 | 16 | 17 | 16 | 12 | 18 | 17 | 18 | 18 | 18 | 53 | 53 | 51 | 30 | 30 | 30 | 30 | 14 | 16 | 16 | 15 | 16 | 5 | 15 | 16 | 15 | 15 | 14 | 16 |
| [TGF-β2] | 18 | 18 | 17 | 16 | 17 | 16 | 12 | 18 | 17 | 18 | 18 | 18 | 53 | 53 | 51 | 30 | 30 | 30 | 30 | 14 | 16 | 16 | 15 | 16 | 5 | 15 | 16 | 15 | 15 | 14 | 16 |
| [TGF-β3] | 16 | 16 | 15 | 14 | 15 | 14 | 10 | 18 | 17 | 18 | 18 | 18 | 51 | 51 | 51 | 28 | 28 | 28 | 28 | 14 | 16 | 16 | 15 | 16 | 5 | 15 | 16 | 15 | 15 | 14 | 16 |
| [MMP-3] | 12 | 12 | 11 | 11 | 11 | 10 | 8 | 16 | 15 | 16 | 16 | 16 | 30 | 30 | 28 | 30 | 30 | 30 | 30 | | | | | | | | | | | | |
| [MMP-9] | 12 | 12 | 11 | 11 | 11 | 10 | 8 | 16 | 15 | 16 | 16 | 16 | 30 | 30 | 28 | 30 | 30 | 30 | 30 | | | | | | | | | | | | |
| [MMP-13] | 12 | 12 | 11 | 11 | 11 | 10 | 8 | 16 | 15 | 16 | 16 | 16 | 30 | 30 | 28 | 30 | 30 | 30 | 30 | | | | | | | | | | | | |
| [TIMP-1] | 12 | 12 | 11 | 11 | 11 | 10 | 8 | 16 | 15 | 16 | 16 | 16 | 30 | 30 | 28 | 30 | 30 | 30 | 30 | | | | | | | | | | | | |
| *Col1* | | | | | | | | | | | | | 14 | 14 | 14 | | | | | 35 | 34 | 35 | 33 | 34 | 19 | 33 | 34 | 35 | 33 | 33 | 34 |
| *Col3* | | | | | | | | | | | | | 16 | 16 | 16 | | | | | 34 | 36 | 36 | 34 | 35 | 19 | 34 | 35 | 35 | 34 | 34 | 35 |
| *Bgn* | | | | | | | | | | | | | 16 | 16 | 16 | | | | | 35 | 36 | 37 | 35 | 36 | 19 | 35 | 36 | 36 | 35 | 35 | 36 |
| *Fmod* | | | | | | | | | | | | | 15 | 15 | 15 | | | | | 33 | 34 | 35 | 35 | 34 | 19 | 33 | 34 | 34 | 33 | 33 | 35 |
| *Dcn* | | | | | | | | | | | | | 16 | 16 | 16 | | | | | 34 | 35 | 36 | 34 | 36 | 18 | 35 | 36 | 35 | 34 | 34 | 35 |
| *Mmp1* | | | | | | | | | | | | | 5 | 5 | 5 | | | | | 19 | 19 | 19 | 19 | 18 | 19 | 18 | 18 | 19 | 19 | 18 | 19 |
| *Mmp3* | | | | | | | | | | | | | 15 | 15 | 15 | | | | | 33 | 34 | 35 | 33 | 35 | 18 | 35 | 35 | 34 | 34 | 33 | 34 |
| *Mmp13* | | | | | | | | | | | | | 16 | 16 | 16 | | | | | 34 | 35 | 36 | 34 | 36 | 18 | 35 | 36 | 35 | 34 | 34 | 35 |
| *Tgfb1* | | | | | | | | | | | | | 15 | 15 | 15 | | | | | 35 | 35 | 36 | 34 | 35 | 19 | 34 | 35 | 36 | 34 | 34 | 35 |
| *Il6* | | | | | | | | | | | | | 15 | 15 | 15 | | | | | 33 | 34 | 35 | 33 | 34 | 19 | 34 | 34 | 34 | 35 | 33 | 34 |
| *Tnfa1* | | | | | | | | | | | | | 14 | 14 | 14 | | | | | 33 | 34 | 35 | 33 | 34 | 18 | 33 | 34 | 34 | 33 | 35 | 34 |
| *Casp3* | | | | | | | | | | | | | 16 | 16 | 16 | | | | | 34 | 35 | 36 | 35 | 35 | 19 | 34 | 35 | 35 | 34 | 34 | 36 |
